# Reusable modular architecture enables flexible cognitive operations in the mouse brain and artificial recurrent networks

**DOI:** 10.1101/2025.08.13.670128

**Authors:** Yuma Osako, Greggory R. Heller, Sofie Ährlund-Richter, Timothy J. Buschman, Mriganka Sur

## Abstract

Complex behaviors are thought to be built by combining simpler cognitive components. Computational modeling has shown that artificial neural networks can perform a variety of tasks by flexibly combining functional modules, each specialized for a specific computation, to construct a complex task. However, it is unknown whether reusable modular networks are found in the brain. Here, we show that mice performing a delayed match-to-sample with delayed report (DMS-dr) task reuse neuronal subspaces that were specialized for stimulus processing and memory maintenance. These subspaces were reused during the task to represent new stimulus inputs and different types of memories, respectively. Clustering analyses showed each subspace was supported by a distinct cluster of neurons in prefrontal cortex and parietal cortex. Studying artificial recurrent networks constrained to neural data found silencing specific clusters disrupted specific computations, consistent with a modular and reusable organization. Altogether, our findings show that the brain can flexibly reuse computational components to perform a complex cognitive task.

## Main text

Real-world behavior in complex environments requires the brain to flexibly coordinate multiple cognitive operations, such as processing sensory inputs, maintaining information in memory, and generating decisions^1–3^. Theoretical models have posited that each of these operations can be performed in independent modules and then reused in different contexts or at different times within the same task^4,5^. This reusability and modularity is thought to allow existing resources to be utilized and combined in different ways, allowing the brain to perform a wide array of cognitive operations and flexibly adapt to new tasks^4–7^. Importantly, reusing neural computations reduces the need to represent all possible combinations of information processing for every task, thereby conserving neuronal resources and achieving the brain’s capacity for generalization. However, despite its theoretical significance, the neuronal circuit mechanisms underlying such reuse of cognitive operations remain largely unknown.

Previous studies in humans^8–11^, non-human primates^12–15^, and rodents^16,17^ have found that task-relevant information is encoded in an abstract, compressed representation that can be reused across contexts. However, these findings primarily focused on the reuse of a specific type of information across different contexts, e.g., reusing the representation of a sensory stimulus for different tasks^9,11,12,14,17^. We refer to this form of reusability as representational reuse, defined here as the repeated engagement of neural circuits to encode the same type of information across different contexts. In contrast, it remains unknown whether the same neural circuit can be repurposed to perform the same computation on different types of information at different times, e.g., maintaining a memory ^18–21^ of a sensory stimulus and then maintaining a memory of an upcoming response (Fig.1a). We refer to this as computational reuse, operationally defined as the engagement of the same neuronal circuit to perform the same type of computation across different task epochs, even when the specific content of the information being processed changes (Supplementary Notes). Addressing this question is critical for understanding the hierarchical level at which the brain reuses neural circuits - whether reuse occurs at the level of representations or computations - thereby providing essential insight into the hierarchy of reusability in neural circuits. Furthermore, while modeling studies suggest that reusable computations are implemented by distinct subpopulations of neurons^4,5^, it is unknown whether this is true in the brain (Fig.1b). Evidence from previous experimental studies has been divided, with evidence that neurons exhibit random mixed selectivity that contributes to multiple task variables^22,23^ and evidence that neurons form distinct clusters, each dedicated to a specific computational function ^24,25^.

**Fig.1.**
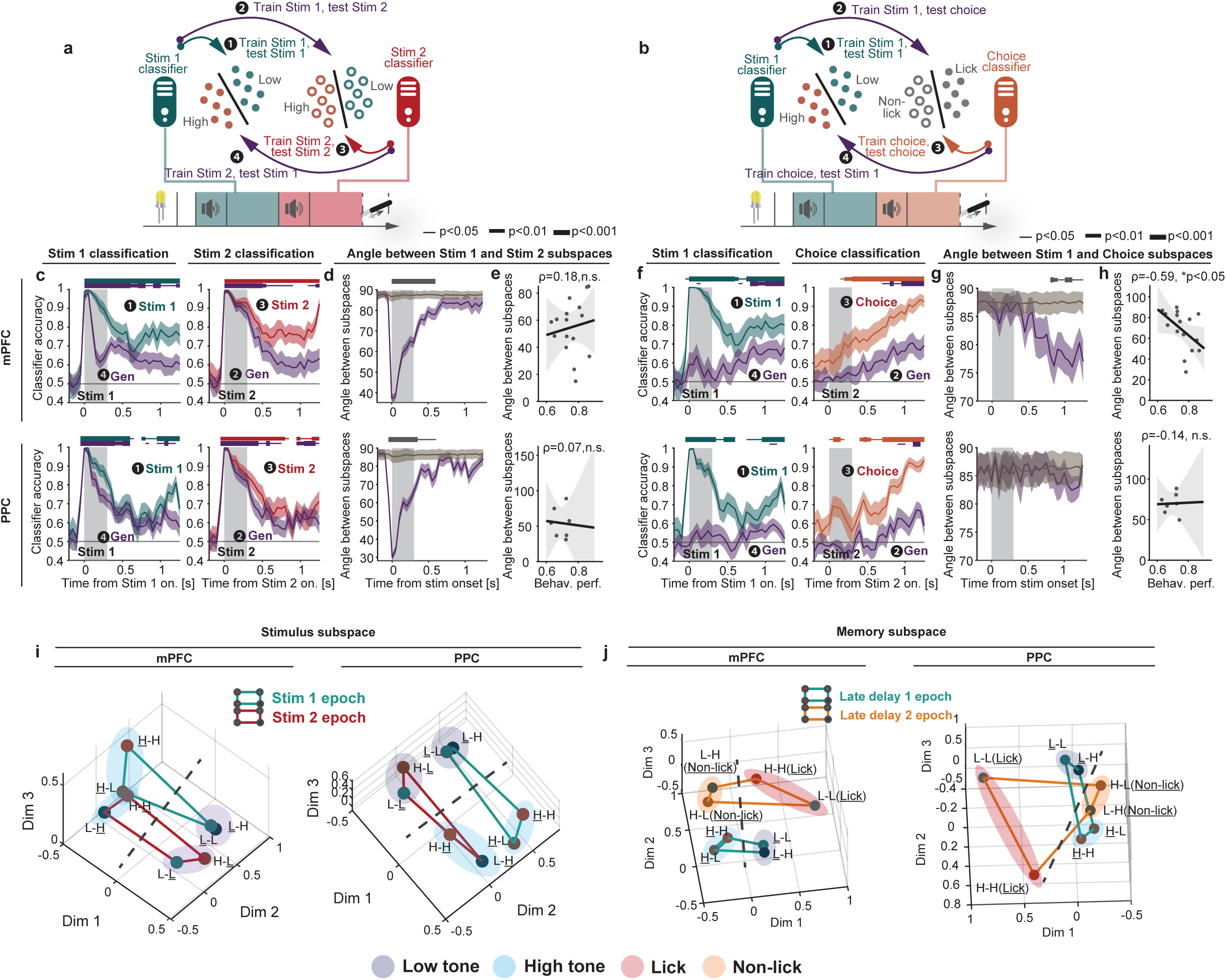
Behavioral performance and neuronal recordings during the DMS-dr task. **(a)** Opposing hypotheses for the neuronal representation of memory content in activity space. One hypothesis (left) posits that distinct information, such as stimuli or choice, is encoded in the different subspaces in neural activity space. Alternatively (right), a shared representational subspace might be utilized across different kinds of information. **(b)** Opposing hypotheses for the functional organization within neuronal populations. One hypothesis (left) suggests that neurons exhibit random mixed selectivity for information processes (e.g., stimulus processing, memory maintenance), with no clear functional clustering. The alternative hypothesis (right) proposes that neuronal selectivity is organized into functionally distinct clusters, where separate subpopulations of neurons are specialized for specific processes or information types. **(c)** Timeline and trial types for the DMS-dr task. Licking was assessed during a 1s response window. **(d)** Experimental setup for the active (left) and passive (right) blocks. **(e)** Correct rate, d-prime, hit rate, and false alarm (FA) rate for all mice (N = 53 sessions, 5 mice). Bar represents mean, and error bar represents SEM. **(f)** Electrophysiological recordings using a high-density probe and neuronal activity of example neurons. Top: spike raster. Bottom: peri-stimulus time histogram (PSTH), with colors representing four different trial types. All firing rates were shown as mean ± SEM. **(g)** Neuronal responses for all neurons in mPFC and PPC to the different trials. Neurons are sorted in descending order of their mean firing rates during the stimulus 1 presentation for H-H trials. **(h)** Proportion of neurons encoding Stim 1, Stim 2 and choice across different task epochs for mPFC and PPC. Bar represents mean, and error bar represents binomial confidence intervals. N=586 and 241 neurons from the mPFC and PPC, respectively.

To address these questions, we developed a delayed match-to-sample with delayed report (DMS-dr) task in mice. In this task, mice were presented with a sequence of auditory stimuli: first a high or low tone (Stim 1) and then, after a memory delay, a second high or low tone (Stim 2). After a second memory delay, the mice were required to report whether the two stimuli matched (or not) by licking (or not licking). This task required the mice to process stimuli during two stimulus presentations (Stim 1 and Stim 2) and and hold task-relevant information over two memory delay periods. We performed electrophysiological recordings to examine neural dynamics in the medial prefrontal cortex (mPFC) and posterior parietal cortex (PPC) during the task. Our findings revealed that different subspaces of neural activity were associated with sensory processing and memory maintenance. These subspaces generalized across time periods. The same stimulus subspace was used to process Stim 1 and Stim 2, reflecting representational reuse and the same memory subspace maintained stimulus information and motor response information in the first and second memory delays, respectively, reflecting computational reuse. Furthermore, sensory processing and memory computations were supported by distinct functional clusters of neurons. When these clusters were inhibited in recurrent neuronal networks (RNNs) trained to reproduce the mouse’s neural activity, the network showed specific deficits in either sensory or memory processes, highlighting the modular architecture that supports flexible cognitive behavior.

## Results

### A task that requires mice to process stimuli and maintain memories twice

We trained mice to perform a delayed matched to sample, delayed response (DMS-dr) task (fig.1c-d). In this task, water-deprived, head-fixed mice were trained to compare two auditory stimuli to determine whether they were the same or different. The stimuli were pure-tone 0.3s auditory stimuli with either a high (H, 14kHz) or low (L, 3kHz) frequency. The first stimulus (Stim 1) and second stimulus (Stim 2) were separated by a 1s memory delay (Delay 1). Then, after the second stimulus, there was another 1s memory delay (Delay 2) before the animal reported whether the two stimuli matched, by licking a reward spout, or did not match, by refraining from licking. Correct licks (hits) were rewarded with water, while incorrect licks to non-match stimuli (false alarm) resulted in an air-puff and a 7s-timeout (Fig.1c). Trials in which the mice refrained from licking after matched stimuli (miss) or after non-match stimuli (correct reject) were not reinforced.

Mice performed the task well, with an accuracy of 76±7%, a discriminability index (d’) of 1.6±0.6, a hit rate of 0.82±0.13, and a false alarm rate of 0.30±0.12 (Fig.1e, mean ± SD, Extended Data Fig.1a-b). Behavioral performance was above chance for all stimulus pairs (Extended Data Fig.1c) and slightly improved after correct trials compared to error trials (Extended Data Fig.1d). To further assess the factors influencing behavioral choice, we fitted a logistic regression model to predict the animal’s response based on the current stimuli and previous trial information (e.g., correctness, stimulus, choice, false alarm, and hit). The model performance was cross-validated (Extended Data Fig.1e, Methods) and showed that mice made decisions primarily based on the current stimuli, with little influence of previous trial information (Extended Data Fig.1f). The reaction time was 138.8 ± 16 ms (Supplementary Fig.1a, mean ± SD), which is faster than that reported in previous studies using similar DMS tasks^26,27^. This suggests the mice already transformed their choice into a motor plan during the delay 2 epoch, prior to the lick port being available (Supplementary Notes, Supplementary Fig.1b-c). Together, these results demonstrate that the mice processed the stimuli and maintained this information in working memory to guide behavior.

To understand how neural representations change in different contexts, mice also performed a block of trials in which they passively listened to the sequence of stimulus (Fig.1d). This block always occurred after the DMS-dr task, and was identical to it, except that the lick spout remained retracted so that the mice could not respond (Supplemental Video 1).

Given the established role of the mPFC and PPC in stimulus processing^28^ and memory maintenance^29,30^, we first tested whether these regions were causally required for the DMS-dr task. We optogenetically inactivated the CaMKII^+^ neuronal activity of the mPFC or PPC during six different epochs using a soma-targeted anion channelrhodopsin (stGtACR2) (Extended Data Fig.1g-h). Inactivation of these areas led to (1) a general decreased correct rate and d-prime across nearly all inhibition epochs, except during the early-delay 1 epoch following PPC inhibition; (2) a decreased hit rate during the stimulus and delay 1 epochs, but an increased hit rate during the delay 2 epoch; and (3) a paradoxical increased false alarm rate across all epochs, except during early delay 1 in both mPFC and PPC, and during late delay 2 in PPC (Extended Data Fig.1i). These results suggest that mPFC and PPC neurons are engaged and are necessary across various epochs of the task.

### Neuronal subspaces are shared across different epochs

To understand how the mPFC and PPC were engaged in the task, we performed dense silicon electrode recordings (Neuropixels probes^31^) from mPFC (515 neurons, n=6 mice) and PPC (241 neurons, n=4 mice) simultaneously (Fig. 1f, left, supplemental table 1). Neurons in both regions represented task-relevant variables, including the identity of the first and second stimulus and the animal’s behavioral response. This information was represented in single neurons (Figs.1f-h and Extended Data Fig.2) and in the neural population (Extended Data Fig.3a-h). To quantify information about these task variables without sampling bias, we trained logistic regression classifiers using all neurons pooled across experiments (“pseudo-population^32^”) to decode stimulus (high/low-tone) and choice (lick/non-lick) (Fig.2a-b). Classifiers trained on mPFC and PPC neural activity accurately decoded both stimulus and choice information (Fig.2a; performance was measured on cross-validation trials). As expected, classifiers for stimulus 2 and choice performed at chance level prior to the presentation of the stimulus 2, indicating that this information was not represented by the neural population until after the stimulus 2 (Fig.2a). To measure the stability of the population code over time, we used trained classifiers at each moment in time and tested their ability to classify at other moments of time (‘cross-temporal classification’, Fig.2b; Methods). To quantify the change in representation, we calculated the angle between the classifiers that represented stimulus 1 during the Stim 1 and Delay 1 epochs (Fig.2c; Methods). In the mPFC, these subspaces were nearly orthogonal, whereas in the PPC they exhibited a partial overlap. These analyses suggest that the population code dynamically shifted immediately following the stimulus presentation (0-0.5s from stimulus 1) into the memory delay (0.8-1.3s from stimulus 1), suggesting that mice utilize different population subspaces during these epochs ^33,34^.

**Fig.2.**
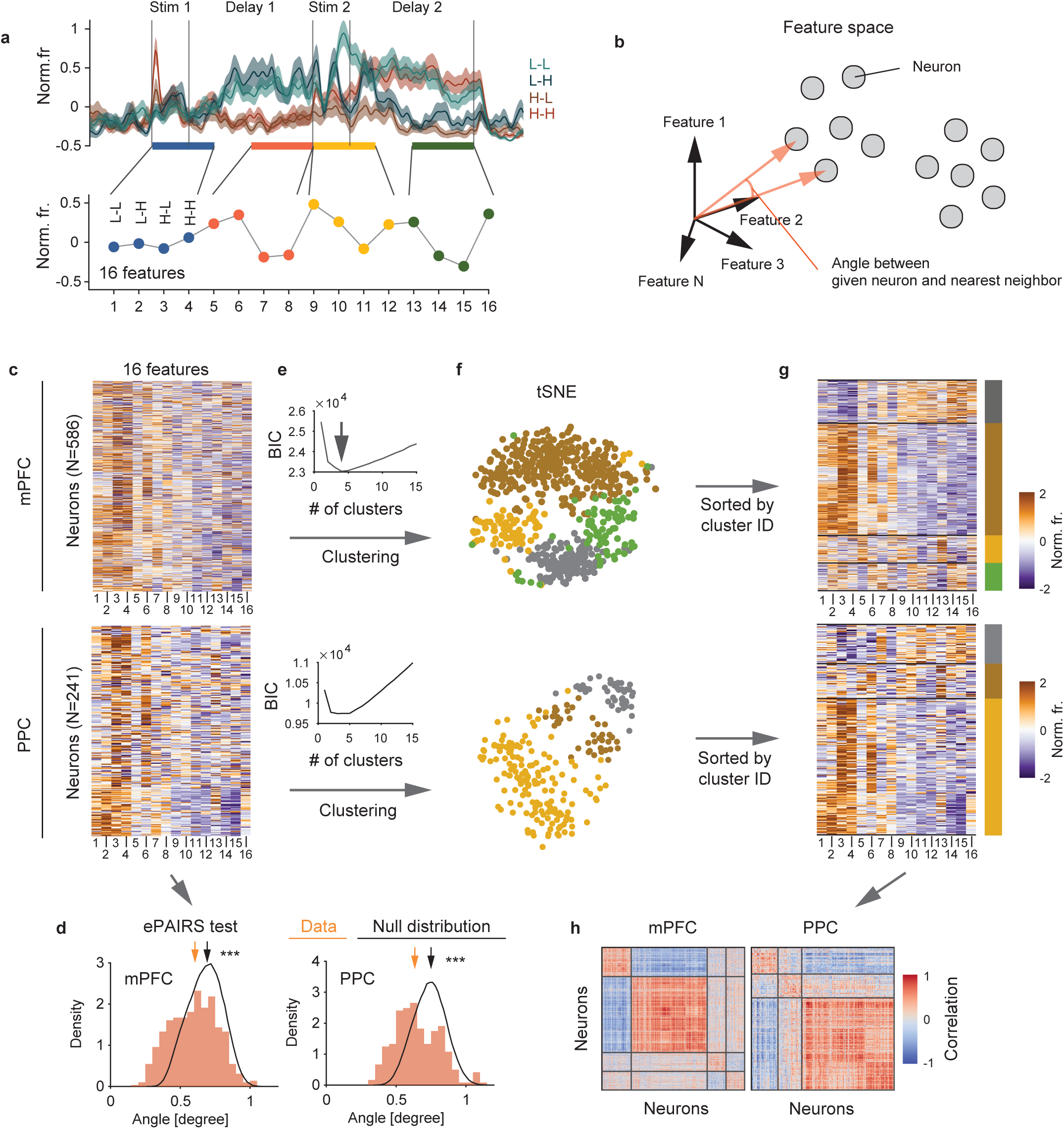
Neural subspaces for stimulus and memory maintenance are orthogonal. **(a)** Time course of classifier accuracy for stimulus 1, stimulus 2, and choice information in the mPFC and PPC. Horizontal bars (top of each plot) indicate above-chance classification (P < 0.05 and 0.01 for thin and thick lines, respectively; one-sided uncorrected bootstrap test). All classifier accuracies were shown as mean ± SD. N = 100 bootstraps. **(b)** Cross-temporal classifier accuracy for stimulus 1, stimulus 2, and choice information in the mPFC and PPC. N = 100 bootstraps. **(c)** The angle between subspaces for stimulus 1 trained during stimulus and late-delay 1 periods. No significance between empirical and shuffled data in the mPFC indicates that these subspaces are orthogonal to each other. One-sided bootstrap test. **p < 0.01. N = 100 bootstraps.

These results suggest that there are dedicated subspaces of neural activity that perform specific cognitive computations – one for processing sensory stimuli and one for maintaining information in memory. If this is true, then the same population representation could be re-used, either at different times during the task or to encode different types of information. To test this hypothesis, we trained logistic regression classifiers to decode stimulus identity (low, L, vs high, H, tones) or choice (lick vs non-lick; i.e., (L-L + H-H) vs (L-H + H-L) trials) from the neuronal population activity and quantified how well these classifiers generalized across conditions and epochs^9,12,13^ (Fig.3a-b, purple arrow, ❷ and ❹). In both mPFC and PPC, classifiers trained to decode the identity of the stimulus during one stimulus epoch generalized this discriminability to the other stimulus epoch ^14^ (Fig.3c, purple ❷ and ❹, p<0.05, bootstrap test, Extended data Fig.4a). In other words, a classifier trained on Stim 1 decoded Stim 2, and vice versa (Fig.3c, Stim 2 classifications, purple). To evaluate if the stimulus subspaces overlapped, we quantified the angle between Stim 1 and 2 period classifiers at each timepoint and found the subspaces were significantly more aligned than expected by chance but gradually became orthogonal during the delay period (Fig.3d, Extended data Fig.4b-c for all combinations of time points). The angle between Stim 1 and Stim 2 subspaces was not correlated with the behavioral performance of animals (Fig.3e, Rho = 0.182 and p = 0.47 for mPFC, and Rho = 0.071 and p = 0.90 for PPC, permutation test). We additionally found that the shared sensory subspace was also engaged during the passive block (Extended Data Fig.4d-f, p<0.05, bootstrap test), indicating that it generalizes across contexts. Together, as expected, these cross-condition classification analyses suggest that a common neural circuit is used to represent the identity of a stimulus, regardless of the time during the task and even during the passive block of trials. This form of ‘representational reuse’ allows stimulus processing to generalize across contexts.

**Fig.3.**
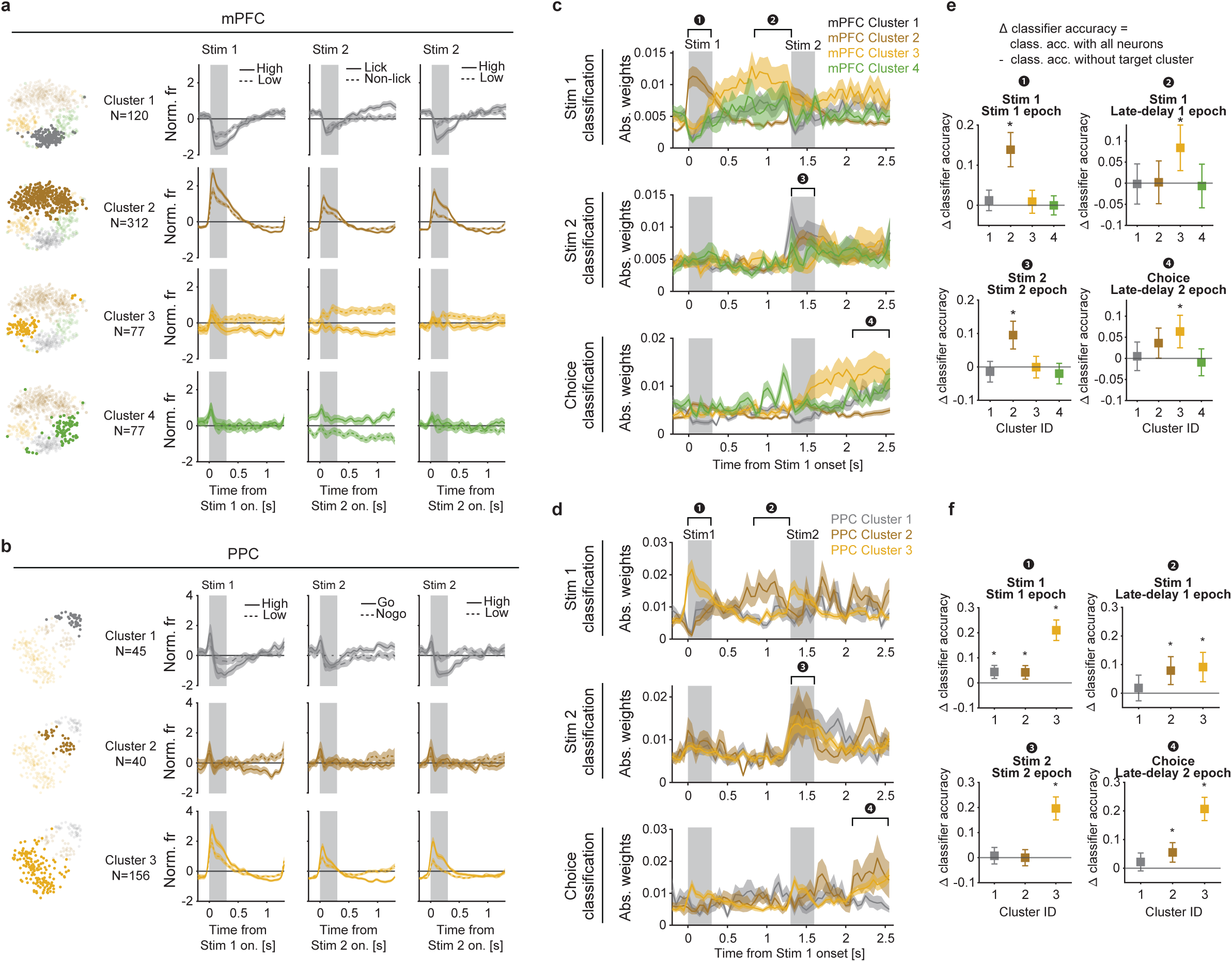
Neural subspaces are shared during the task. **(a-b)** Schematic of classifiers used to quantify stimulus and choice information. Stimulus 1, stimulus 2, and choice information are evaluated during [-0.2s 1.3s] from stimulus 1 or 2 onset. Cross-conditional classification was performed by decoding stimulus 1 (or 2) using a classifier trained on stimulus 2 (or 1) (**a, 2** and **4**) or by decoding stimulus 1 (or choice) using a classifier trained on choice (or stimulus 1) (**b, 2** and **4**). **(c)**. Time course of classifier accuracy for stimulus 1 and 2 information in each region. Horizontal bars (top of each plot) indicate above-chance classification (*p* < 0.05 and 0.01 for thin and thick lines, respectively). N = 100 bootstraps. **(d)** Time course of the angle between stimulus1 and 2 subspaces for each region. Horizontal bars (top of each plot) indicate above-chance classification (*p* < 0.05 and 0.01 for thin and thick lines, respectively). All classifier accuracies were shown as mean ± SD. N = 100 bootstraps. **(e)** Angle between stimulus 1 and 2 subspaces as a function of behavioral performance. Each dot represents a single session (N=16 or 7 sessions for mPFC and PPC, respectively). Shaded area shows 95% confidence interval. Spearman’s rank-order correlation coefficient was used. **(f)** Same as (c), but for stimulus 1 and choice information in each region. N = 100 bootstraps. **(g)** Same as (d), but for angle between stimulus1 and choice subspaces for each region. N = 100 bootstraps. **(h)** Same as (e), but for angle between stimulus 1 and choice subspaces as a function of behavioral performance. Each dot represents a single session (N=16 or 7 sessions for mPFC and PPC, respectively). **(i-j)**. Representations of different trials during the stimulus epoch (0-0.5s and 1.3-1.8s from stimulus 1 onset) (**i**) and the late-delay epoch (0.8-1.3s and 2.1-2.6s from stimulus 1 onset) (**j**) for each region. Lines and shading represent mean ± SD for **c**, **d**, **f**, and **g**. For **e** and **h**, lines are fitted with least-squares linear regression. n.s. non-significant, *p<0.05, Two-sided permutation test. Shading denotes the 95% confidence interval of the regression. For **c**, **d**, **f**, and **g**, one-sided uncorrected bootstrap tests were performed for statistical test.

In the DMS-dr task, mice were required to hold two distinct types of information during the first and second memory delays: during the first memory delay, they had to remember the recent sensory stimulus, and during the second memory delay, they remembered the upcoming behavioral response (Fig.1a). Despite these being different types of information, we were curious as to whether the same subspace was used to maintain both memory representations. In mPFC, classifiers trained to decode stimulus identity during delay 1 significantly decoded choice during delay 2, and vice versa (Fig.3f, purple ❷, p<0.05, bootstrap test, Extended Data Fig.5a-c). In other words, a classifier trained to decode the identity of Stim 1 during the first memory delay was able to decode the animal’s upcoming choice during the second memory delay, and vice versa (Fig.3f, choice classifications, purple ❹). These shared subspaces were not attributable to a selectivity bias towards the tone (Extended Data Fig.5d) or due to the licking action itself (Extended Data Fig.5e) and the same effect was seen in single-sessions (Extended Data Fig.5f). Note that the identity of the Stim 1 and choice were not correlated because we balanced trial numbers across conditions (see Methods). Therefore, the observed generalization of the memory subspace cannot be accounted for by a shared representation of Stim 1 and Stim 2 during the delay periods (Supplementary Fig. 2). The angle between the subspace representing the memory of the first stimulus and the memory of the upcoming choice decreased throughout the delay period in the mPFC (Fig.3g), suggesting the two representations became more similar over time. This is in contrast to the angle between the memory subspace and the shared stimulus subspace (Fig.3d). In general, the shared subspace was stronger in mPFC compared to PPC (Fig.3g). Similar results were also found using a linear regression model (Supplementary Fig.3). Together, these results suggest that, in mPFC, there was a dedicated subspace of neural activity that maintained memories, regardless of the exact type of information being maintained (e.g., both the memory of the first stimulus and the memory of the upcoming choice). This form of ‘computational reuse’ would allow a neural circuit to be specialized for a specific cognitive computation that can then be generalized across different contexts.

The engagement of this shared subspace was related to the task. In the mPFC, but not in PPC, lower angles between classifiers, and thus greater subspace overlap, was correlated with better behavioral performance (Fig.3h, Rho = -0.59 and p = 0.02 for mPFC, and Rho = -0.14 and p = 0.78 for PPC, permutation test, Extended Data Fig.5f, bottom). This correlation emerged during the late-delay period (Extended Data Fig.5g). Furthermore, the same subspace was not observed during the passive block; the classifier trained to decode choice (in delay 2) during the DMS-dr task session did not decode the memory of the stimulus in the passive block (Extended Data Fig.5h-i). These findings suggest that the shared subspace for memory maintenance is engaged in a task-specific manner.

Finally, we were interested in whether the shared subspaces preserved the geometry of representations within the neural population. So, we investigated how the neural population represents different trial types by projecting the neural representation of four different trial types (stimulus 1 and 2 pair: Low-Low, Low-High, High-Low, High-High) in the mPFC and PPC into a reduced dimensional space specific to stimulus and late-delay epochs (Method). Projection in the stimulus subspace revealed relatively parallel and aligned coding direction of stimuli (high or low tones) (Fig.3i). This geometric structure permits a single decision boundary (Fig.3i, dotted line) to decode stimulus identity in both stimulus 1 and 2 epochs, suggesting that this boundary is reusable across these stimulus presentations as expected ^11–13^. Projection in the memory maintenance subspace also revealed that a single decision boundary can be used to differentiate stimulus identity and choice (lick or non-lick). However, this boundary was more obvious in mPFC compared to PPC. These geometries emerged only at the corresponding times during the trials (Extended Data Fig.6a-b, Task, stimulus subspace: 0-0.5s and 1.3-1.8s, Extended Data Fig.6c-d memory maintenance subspace: 0.8-1.3s and 2.1-2.6s) and only the stimulus subspace exhibited the same geometry in the passive block (Extended Data Fig.6a-d, Passive).

Interestingly, the sensory subspace and memory subspace in mPFC were aligned such that the coding direction from Low tone to High tone was aligned with the coding direction from Go to No-go (Fig.3j). While seeming to be an arbitrary association, we wanted to test whether training history might explain this alignment. To this end, we pre-trained RNNs on a restricted subset of trial types, either Low–Low and Low–High or High–High and High–Low trials (Extended Data Fig.7a-b). After this pre-training phase, the networks were trained on the full DMS-dr task containing all tone combinations. We found that the final coding direction in the memory maintenance subspace was biased by the pre-training history – when Low-Low and Low-High were associated with lick and no-lick, respectively, we found Low/High were aligned with lick/no-lick (and vice versa for High-Low and High-High, Extended Data Fig.7c-e). This suggests the network began by associating stimulus 2 with the motor response and then this association biased the direction of encoding in the shared memory subspace. Importantly, when RNNs were trained without pretraining the association between stimulus and choice was arbitrary and random for each individual RNN. Yet, these networks exhibited the same form of computational reuse (Extended Data Fig.7e). Notably, mice were trained using the Low–Low and Low–High trial combinations during the early phase of training (same as 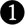 RNN in Extended Data Fig.7a) and exhibited results similar to the RNN simulations (Extended Data Fig.7f-g). Moreover, the degree of alignment between coding directions was not correlated with the duration of pre-training or DMS-dr training (Extended Data Fig.7h), but instead was significantly correlated with behavioral performance during pre-training (Extended Data Fig.7i, rho=0.6545, p = 0.0260, permutation test). Together, these results suggest that training history biases coding direction within the memory maintenance subspace.

Altogether, these results suggest that, as expected, the neural subspace for stimulus representation is dynamically reused at appropriate times in both the mPFC and PPC. In contrast, the subspace for memory maintenance exhibits reusability especially in the mPFC. Notably, memory maintenance subspaces are reused even when the information they encode is different, suggesting that this subspace serves a specific cognitive computation rather than merely reflecting the physical similarity of stimuli. The importance of this subspace is highlighted by the fact that the animals’ behavioral performance was greatest when the memory maintenance subspace in the mPFC was more shared.

### Single neuron clusters represent distinct neuronal dynamics

Thus far, we have shown that subspaces are shared at appropriate times during the task, supporting the concept of reusable cognitive operations. These operations could be interpreted either in terms of collective dynamics from all neurons (Fig.1b, left, Random mixed selective)^22,23,35^ or as subpopulations of distinct neurons^24,25^ (Fig.1b, right, Clustered). To distinguish between these hypotheses, we employed two approaches. First, we examined the response profile of individual neurons during the task. A response vector was created for each neuron by concatenating its responses across the four trial types (L-L, L-H, H-L, H-H) during the stimulus and late-delay epochs for both the first and second stimuli (Fig.4a, 4 trial types × 4 epochs = 16 dimensions). This response vector from all neurons allowed us to examine the distribution of neurons within the feature space. A neuronal population that is randomly mixed within the representational space would have uniformly distributed response vectors around the origin. To test this, we applied the ePAIRS test (Fig.4b, Methods)^22,24^ to compare the empirical distribution with a randomized shuffle corresponding to a multivariate Gaussian (Fig.4c-d). Both mPFC and PPC populations had non-uniform distributions, significantly different from the artificial uniform distribution (Fig.4d, p<0.001, ePAIRS test, Methods). This indicates that individual neurons in these regions do not uniformly occupy the space of all possible representations but instead have functional specializations. This non-random distribution was not observed when using neuronal activity during pre-stimulus period (Extended Data Fig.8a), suggesting functional specialization is associated with information processing.

**Fig.4.**
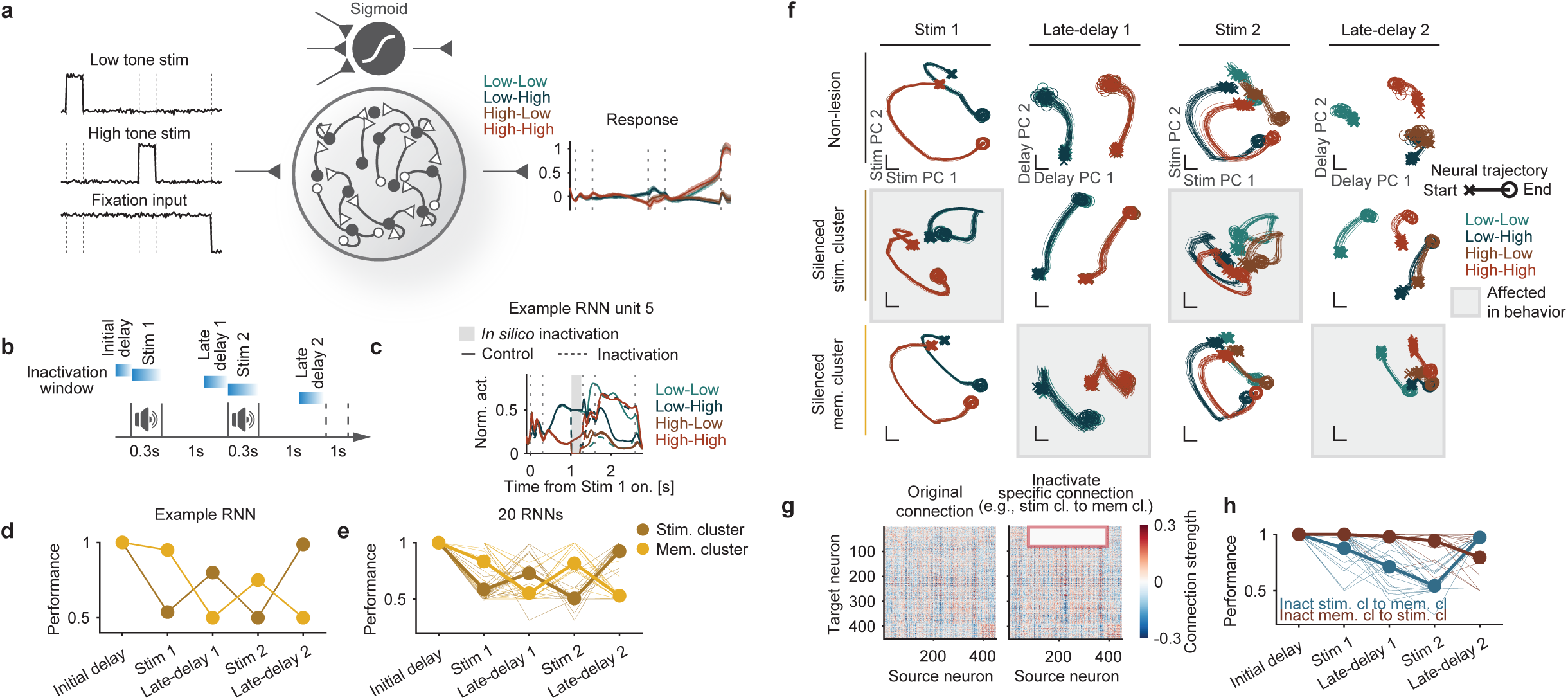
mPFC and PPC neurons form discrete clusters. **(a)** Top: neural response of an example neuron across all four trial types. Bottom: response vector calculated as the mean response during each epoch (0-0.5s, 0.8-1.3s, 1.3-1.8s, and 2.1-2.6s from stimulus 1 onset). Neuronal activities are shown as mean ± SEM. **(b)** A 16-dimensional vector represents each neuron’s activity to each feature. For neuronal populations with random mixed selectivity, vectors are uniformly distributed around the origin, which can be quantified by angles between nearest neighbors (ePAIRS). **(c)** Unsorted response vectors of all neurons in the mPFC. N = 586, 241 neurons in mPFC and PPC, respectively. **(d)** Distribution of angles between each point and its nearest neighbor in the feature space compared to that of a matching multivariate Gaussian (null distribution, black line) for each brain region. \*\*\**p*<0.001, ePAIRS test (two-sided, see Methods). N = 586, 241 neurons for mPFC and PPC. N = 500, 500 bootstraps for null distributions for mPFC and PPC, respectively. **(e)** BIC scores used to determine the optimal number of clusters. **(f)** Visualization of the response vectors using t-distributed stochastic neighbor embedding (t-SNE). Each dot represents a single neuron, colored according to its cluster membership. N = 586, 241 neurons for mPFC and PPC. **(g)** Response vectors sorted by their respective cluster membership. N = 586, 241 neurons for mPFC and PPC. **(h)** Sorted correlation matrices revealing strong within-cluster similarity. N = 586, 241 neurons for mPFC and PPC.

Next, we tested whether neurons clustered into functional modules. Fitting a Gaussian mixture model (GMM) to the response vectors (Fig.4c, e-g, Methods) revealed four functional clusters in mPFC (N=120, 312, 77, 77 for cluster 1-4, respectively) and three functional clusters in PPC (N=45, 40, 156 for cluster 1-3, respectively) (Fig.4e; chosen based on minimizing Bayesian information criterion, BIC, scores). For visualization, we used tSNE^36^ to project neural responses into a two-dimensional space (Fig.4f). Consistent with the observed clustering, all clusters showed strong correlation of activity within the cluster (Fig.4h) and significantly greater co-fluctuation among neurons within the same cluster compared to across clusters (except mPFC-cluster 1 and PPC-cluster 2, Extended Data Fig.8b-e). The functional clusters were not distinguished by waveform of neurons or anatomical locations within mPFC and PPC (Extended Data Fig.8f-h), suggesting that functional specialization arises from network dynamics rather than fixed anatomical segregation.

Each cluster represented distinct patterns of neural representation (Fig.5a-b; confirmed with a linear classifier, see Extended Data Fig.8i and Methods for details). mPFC-clusters 1 and 2 distinguished between high and low tones during stimulus 1 and 2 (Fig.5a, mPFC-clusters 1 and 2), while neurons in mPFC-cluster 3 differentiated high/low tones and lick/non-lick behavior during delays (Fig.5a, mPFC-cluster 3). mPFC-cluster 1 also distinguished between lick and non-lick behavior during the second delay, and mPFC-cluster 4 might represent behavioral bias (Fig.5a, cluster 4). In the mPFC, we observed clusters that were selectively engaged in distinct phases of the task, such as those dedicated to processing sensory information during the stimulus periods (Fig.5a, mPFC-cluster 2) or maintaining information during the delays (Fig.5a, mPFC-cluster 3). In contrast, such specialized patterns of activity were less apparent in the PPC. For example, PPC-clusters 1 and 3 represented sensory information during the stimulus periods but also encoded lick/non-lick behavior during the second delay, suggesting a multiplexed coding scheme in PPC ^22,37^.

**Fig.5.**
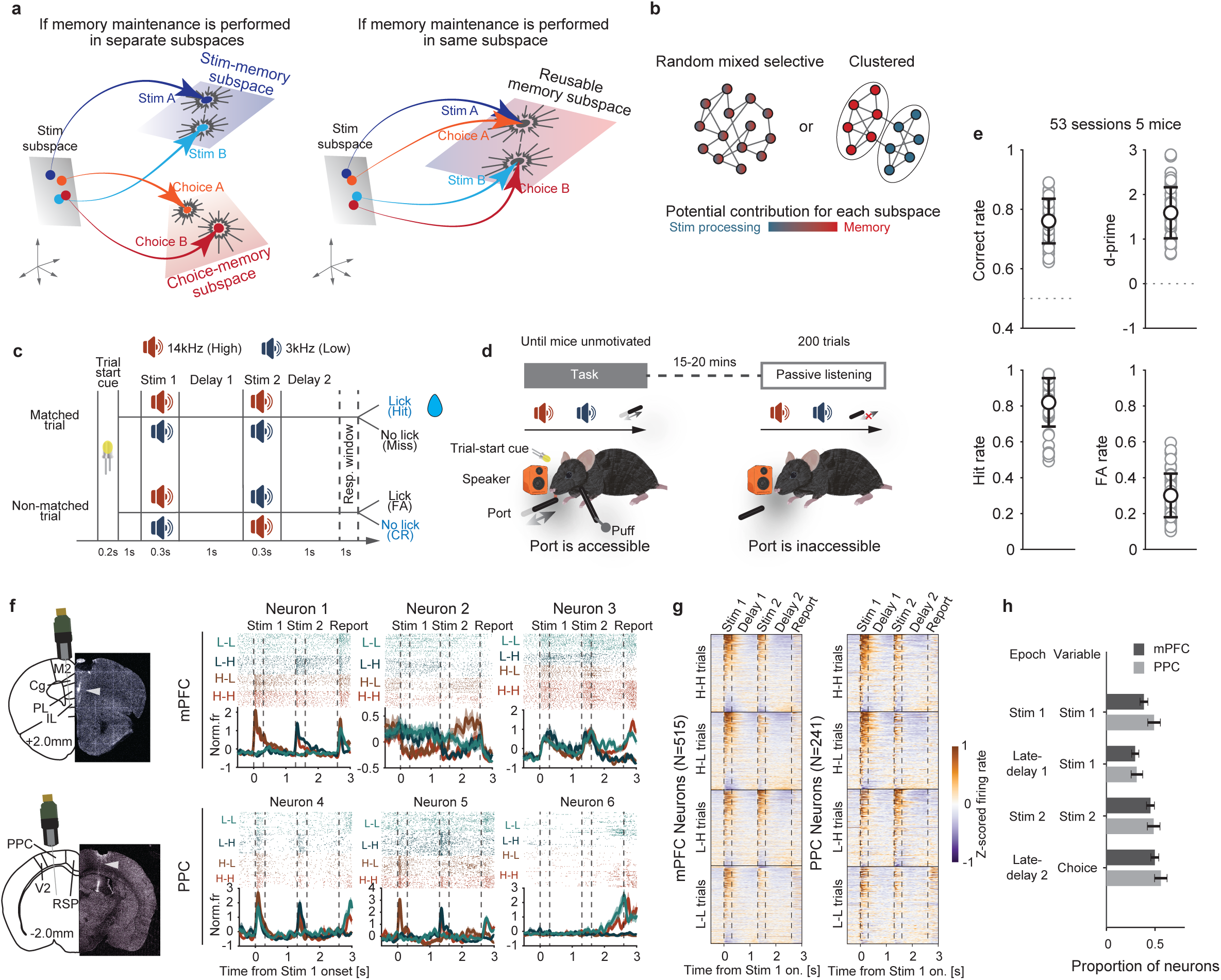
Discrete clusters dominantly contributed to subspaces at different timing. **(a-b)** Average responses of neurons within each cluster. All neuronal activities are shown as mean ± SEM. **(c-d)** Time course of absolute weights for the stimulus 1, stimulus 2, and choice classifiers. All weights are shown as mean ± SEM. Each color corresponds to cluster ID. **(e-f)** Change in classification accuracy for each cluster was computed by subtracting the classification performance computed after removing the target cluster from the performance using all neurons. N = 100 bootstrap. \**p*<0.05. One-sided bootstrap test with Holm-Bonferroni correction. All plots are shown as mean ± 2SD.

Given these responses, we hereafter refer to mPFC-cluster 2 and PPC-cluster 3 as putative stimulus clusters and mPFC-cluster 3 as the putative memory maintenance cluster. To further investigate the contributions of each functional cluster to the stimulus and memory-maintenance subspaces, we calculated the absolute weights of each cluster for the classifiers trained above (Fig.5c and 5d). The putative stimulus clusters, i.e., mPFC-cluster 2 and PPC-cluster 3, contributed most strongly to the sensory classifiers trained to decode the identity of the first stimulus during the stimulus 1 epoch (Fig.5c, brown line in stimulus 1 classification for mPFC. Fig.5d yellow line in stimulus 1 classification for PPC). Moreover, the mPFC-cluster 2 also contributed to the second stimulus classifier during the stimulus 2 epoch (Fig.5c and 5e, brown in stimulus 2 classification). For the memory classifiers, which were trained to decode first stimulus during delay 1 or choice information during delay 2 epochs, the putative memory maintenance cluster, i.e., mPFC-cluster 3 and PPC-cluster 2 and 3, contributed significantly during both delay 1 and delay 2 (Fig.5c-f, yellow in mPFC, brown in PPC). Moreover, mPFC-cluster 3 showed a sustained representation of information across the delay, while PPC-cluster 2 exhibited a ramping pattern leading up to the choice (Supplementary Fig. 1c). We also confirmed that putative excitatory neurons are engaged in both stimulus- and memory-related dynamics (Supplementary Fig.4).

Together, these results suggest that stimulus and memory maintenance clusters in mPFC are engaged by different computations; sensory processing recruits sensory clusters during stimulus epochs while memory processing recruits memory maintenance clusters during delay epochs. In contrast, clusters in the PPC exhibited multiplexed contributions, participating in both stimulus and memory representations across different task epochs.

### Modular lesion effect in an RNN model of mPFC subpopulations

Our data suggest there are reusable neural subspaces for stimulus processing and memory maintenance, each supported by distinct subpopulations of neurons. We next trained RNNs to investigate potential neural circuit mechanisms underlying the modular architecture of stimulus processing and memory maintenance observed in mPFC (Fig.6a). Our aim of this approach was to explore how distinct subpopulations within a network can give rise to flexible, epoch-dependent computations. Specifically, we sought to understand (1) how stimulus- and memory-maintenance subpopulations can interact within a network, and (2) how perturbations to these subpopulations affect task performance and network dynamics. While multiple underlying network architectures can generate similar population trajectories, this approach allows us to test mechanistic hypotheses about neural reusability and modularity beyond descriptive observations. The RNNs reliably reproduced the animals’ behavior and the PSTHs of mPFC neurons (Extended Data Fig.9a-b). We also found that stimulus cluster units received stronger connection weights from stimulus inputs when compared to memory maintenance cluster units. In contrast, memory maintenance cluster units exhibited stronger connections to the output units than stimulus cluster units (Extended Data Fig.9c). Units whose activity was not constrained by neuronal activity in mPFC exhibited heterogenous responses to the task (Extended Data Fig.9d). To assess the functional role of the neuron clusters observed in Figure 5, we silenced each cluster during different epochs of the DMS-dr task (Fig.6b-c), including the initial delay, stimulus 1, late-delay 1, stimulus 2, and late-delay 2 epochs. Task performance was impaired when the mPFC-cluster 2 stimulus cluster was silenced during the stimulus epochs and when the mPFC-cluster 3 memory cluster was silenced during the late-delay epochs (Fig.6d). This effect was consistent across multiple simulations (Fig.6e). Importantly, this modular lesion caused no impairment when the putative stimulus or memory maintenance cluster was not actively engaged. To further understand each cluster’s role in cognition, we overrid the activity of each cluster with activity from a different trial type during specific task epochs (Extended Data Fig.9e). This activity-patching perturbation showed similar modular effects as the in-silico inactivation experiments (Extended Data Fig.9f). Examining the network state revealed that silencing the putative stimulus processing cluster during stimulus presentation (Stim 1 and Stim 2 epochs) significantly impacted state trajectories (Fig.6f, Stim 1 and Stim 2, silenced stim cluster). In contrast, silencing the putative memory maintenance cluster during these periods had minimal impact (Fig.6f, Stim 1 and Stim 2, silenced mem cluster). On the other hand, during late-delay periods, silencing the putative memory maintenance cluster altered state trajectories. Meanwhile, silencing the putative stimulus processing cluster had little effect on state trajectories although the relative positions of each trial type remained unchanged (Fig.6f, Late-delay1 and Late-delay2). Finally, we found that silencing the projection from the mPFC-cluster 2 stimulus cluster to the mPFC-cluster 3 memory maintenance cluster during stimulus epochs impaired behavioral performance (Fig.6g-h), suggesting the putative stimulus processing cluster conveys information to the memory maintenance cluster during stimulus presentation. Altogether, these findings are consistent with a modular organization of stimulus processing and memory maintenance clusters in the network, providing a candidate circuit mechanism for flexible computations during the task.

**Fig.6.**
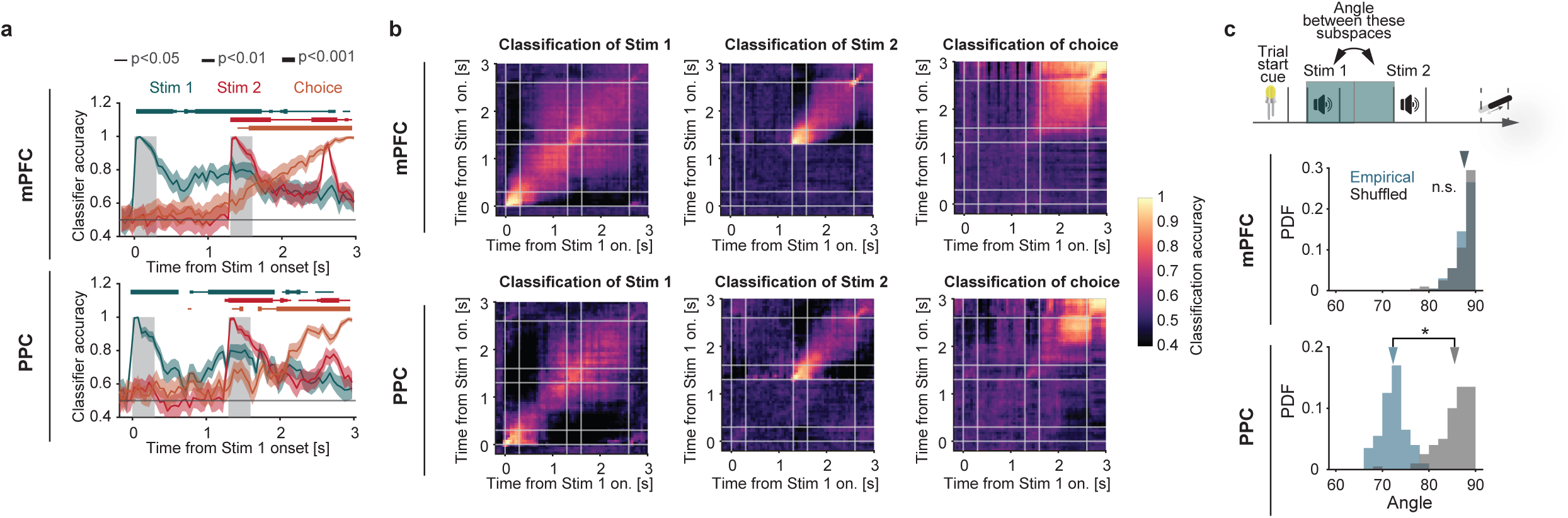
Data-constrained RNN shows modular lesion effects. **(a)** Example of a fully connected RNN architecture. The network receives stimulus and fixation inputs and projects to a fixation output unit (active when a motor response is unwarranted) and a behavioral output unit. All RNN units exhibit non-negative response. RNN containing 77 and 312 units were trained to mimic the neuronal activity of cluster 2 and 3 in the mPFC (Methods). Example RNN outputs are shown as mean ± SEM. **(b)** Schematic illustrating timing of *in-silico* inactivation of RNN clusters. **(c)** Example activity of the putative memory maintenance RNN unit during late-delay1 period, with either the putative stimulus or memory maintenance cluster inactivated. Solid and dashed lines indicate the control and inactivation trials. **(d)** RNN performance after silencing specific clusters during each epoch. Yellow and brown lines represent silences of the putative stimulus and memory maintenance clusters, respectively. **(e)** Summary of RNN performance after silencing clusters. Each line represents a different RNN (N=20 networks). Stimulus epochs include stimulus 1 and 2, while delay epochs include late-delay1 and 2. Error bars indicate mean ± SD. **(f)**. State trajectories (starting from ‘x’ and ending with ‘o’) during the performance of stimulus and late-delay periods (0-0.5s, 0.8-1.3s, 1.3-1.8s, 2.1-2.6s from stimulus onset) in silenced and full network projected into the first two PCs defined by the full network state at the stimulus and late-delay epochs, respectively. Each line represents an individual trial. **(g)** Synaptic weights between units. Inactivation of synaptic connections was simulated by setting connection weights to zero at corresponding time (right, rectangle area). **(h)**. RNN performance after silencing specific connection during each epoch. Blue lines represent silence of the connections from the putative stimulus processing cluster to memory maintenance cluster, and brown lines represent opposite direction. Thin lines represent individual networks.

## Discussion

A combination of analyses of empirical neuronal data and computational simulations suggest the brain uses dedicated computational circuits for performing specific cognitive operations. Mice trained to perform a delayed match-to-sample task with a delayed response, used a ‘sensory’ subspace of neural activity in both mPFC and PPC to process incoming stimuli. The sensory subspace was reused across different contexts^9,12–15,17,18,38^, reflecting its ability to generalize. Moreover, we found that a ‘memory’ subspace in mPFC maintained the memory of task-relevant information. Importantly, the same memory circuit was used to maintain different types of information. In the first memory delay, the memory subspace represented the retrospective memory of the presented stimulus while, in the second memory delay, the memory subspace represented the prospective memory of the upcoming response (Fig. 3f-h, and 3j). This suggests the memory subspace engaged a specific cognitive computation – maintenance of information – that was agnostic as to the contents of the memory itself. Together, our results suggest that mPFC not only generalizes specific information representation across various contexts but can also generalize cognitive operations across different categories of information, reflecting a form of computational reuse. This computational reuse was important for behavior: the degree to which memory maintenance subspaces were shared was significantly correlated with behavioral performance (Fig.3h and Extended Data Fig.5f, bottom). Whether these functional clusters are learned or are pre-existing remains an open question. Previous work suggests that memory maintenance neurons emerge, and their representations stabilize as animals learn the task^39^, which implies that the brain might co-opt existing modules, such as a memory maintenance module, during learning^40^.

It has been debated whether neurons are organized into clusters according to their functional roles^24,25^ or if responses are randomly distributed^22,23^. Recent work has typically found a non-modular architecture in most brain areas^41^. In contrast, we found functional clustering (see also ^24,25^). Neurons in both regions were clustered according to their response patterns during the active task (Fig.4), with specific clusters contributing predominantly to stimulus processing and memory maintenance subspaces, respectively (Fig.5). Moreover, when ablating these clusters in RNNs trained to reproduce our experimentally observed neural dynamics, we found the sensory and memory clusters impaired specific aspects of behavior and disrupted the corresponding functional dynamics of neural activity (Fig.6). Together, our results suggest that there are modules of neurons performing cognitive operations and that these modules can be flexibly reused across time and tasks.

The discrepancy between our results and previous work might stem from differences in the behavioral tasks used. Studies reporting non-modular architectures often utilize tasks such as multisensory decision-making or visual-spatial detection, where the stimulus is directly linked to behavior. By contrast, Dubreuil and Valente et al. reported that networks performing tasks that require flexible input-output mapping exhibit subcluster organization^42^. In line with this idea, the DMS-dr task in our study demands flexible output mapping based on combination of sensory inputs (e.g., the same first stimulus leads to different responses depending on the identity of the second stimulus). This requirement for dynamically integrating multiple inputs might promote the emergence of subcluster organization within the neural network.

Within the memory maintenance subspace, we observed that low and high tones were generalized to correspond with lick and non-lick responses, respectively (Fig.3j). This direction of generalization was consistent across animals and might reflect the sequence of task training (Extended Data Fig.7f-g, 14 out of 15 sessions, 92%). During the early stage of training, the mice were first exposed to low-low and low-high pairings (Extended Data Fig.7f, bottom), where the low tone during stimulus 2 was directly associated with the lick response. Although high-high and high-low pairings were subsequently introduced later during training, we speculate that the earlier training experience promoted an initial association between low tones and reward, causing low tones and lick behavior to become aligned along the same direction in the subspace. This effect was reproduced in RNN simulations (Extended Data Fig.7a-e). These findings suggest that the specific geometric features of memory maintenance representation are sculpted and may be associated with behavioral biases. This highlights a critical insight: rather than simply organizing information in a generalized geometric manner, the brain also encodes subtle, behaviorally relevant details within this generalized framework such as bias.

Our findings suggest that in the mPFC, distinct subpopulations of neurons selectively contribute to subspaces associated with stimulus processing and memory maintenance (Fig.5a, c, e). In contrast, in the PPC, these subspaces were supported by multiple neuronal subpopulations (Fig.5b, d, f). This difference implies that while the mPFC may possess a more strongly modular architecture, the PPC may rely on neuronal populations that exhibit more multiplexed activity patterns. Notably, although neuronal responses in the PPC could still be categorized into clusters based on their tuning properties, these clusters did not map onto specific functional subspaces in a one-to-one manner. This observation is consistent with previous study suggesting more distributed or mixed selectivity in parietal areas^22^. Furthermore, the memory-maintenance subspace in the mPFC exhibited strong reusability across different types of information (Fig.3f-g and 3j). This was weaker in the PPC, suggesting that PPC allocates partially distinct subspaces to working memory representations depending on the content being maintained. In contrast, the mPFC may operate at a more abstract level, providing a content-independent computational resource for sustaining information.

Our RNN simulations, constrained by empirical data, demonstrated the effect of lesioning specific functional modules in the mPFC (Fig.6). These findings align with previous theoretical studies^4,5,19^ reinforcing the concept of modular architecture and its reusability within the mPFC. We note that functional subspaces are not unique to networks with non-negative activation functions. Rather, non-negativity may bias these representations toward axis-aligned implementations, thereby making distinct functional subpopulations more readily identifiable at the single-unit level. Interestingly, silencing synaptic connections from the stimulus processing to the memory maintenance cluster during stimulus periods disrupted network behavior (Fig.6f-g). This suggests that in spite of the modular organization, the recurrent circuitry connecting these modules is necessary and sufficient for driving network dynamics^43–45^. If the modular organization of these clusters in animal brains mirrors the architecture observed in our simulations, time-specific neural manipulations such as holographic perturbation^46–50^ could further uncover the underlying neural mechanisms of such architecture. Future work will be required to test this prediction and further elucidate the modular architecture and its reusability underlying flexible cognitive behavior.

In conclusion, our study provides evidence for clusters of neurons that are dedicated to specific cognitive computations. These computations can be reused across time and tasks. These findings suggest that reusable modular architecture underpins the complexity and flexibility of animal behavior. Furthermore, our results provide important insights into the significance of generalization in information representation, extending our understanding of how the brain supports this crucial cognitive function.

## Acknowledgements

We thank members of the Sur and Buschman labs, especially Yi-Ning Leow, Ruby Lam, Prachi Ojha, Gabrielle T. Drummond, Tatsuya Osaki, and Adel Ardalan for fruitful discussion and advice; and Stefano Panzeri, Yoshihito Saito, Toshitake Asabuki, Kensuke Yoshida, and Tomoya Ohnuki for constructive feedback on this paper. We also thank Arundhati Natesan and Emma Odom for slicing and imaging mice brain.

## Author Contributions

Conceptualization: Y.O., T.J.B.

Methodology: Y.O., G.R.H., S.Ä-R

Software: Y.O.

Validation: Y.O.

Formal analysis: Y.O.

Investigation: Y.O.

Resources: Y.O., M.S.

Data Curation: Y.O.

Writing - Original Draft: Y.O.

Writing - Review & Editing: Y.O., G.R.H., S.Ä-R, T.J.B., M.S.

Visualization: Y.O.

Supervision: T.J.B., M.S.

Project administration: Y.O.

Funding acquisition: Y.O., T.J.B., M.S.

## Methods

### Mice

All procedures conducted in this study were approved by the Massachusetts Institute of Technology’s Animal Care and Use Committee and conformed to the Guide for the Care and Use of Laboratory Animals published by the National Institutes of Health. Male and female mice more than 8 weeks old were used in this study. Mice were housed in a room with reversed light/dark cycle (light off from 09:00 to 21:00) with controlled temperature and ventilation (20– 22 °C; 40–60% humidity). All experiments were performed during the dark period of the cycle.

### Virus

For inactivation experiments, CaMKIIa-driven soma-targeted anion-conducting channelrhodopsin fused to FusionRed (pAAV-CKIIastGtACR2-FusionRed, Addgene, 105669; titre, 1 × 10^13^ viral genomes per ml) was used to express GtACR2 in the soma of excitatory neurons.

### Stereotactic surgeries

Animals were prepared similarly for all surgical procedures. Mice were anaesthetized using isoflurane anaesthesia (3% for induction, 1–1.5% for maintenance) while maintaining a body temperature of 37.5 °C using a heating pad (ATC2000, World Precision Instruments). Mice were given pre-operative slow-release buprenorphine (1 mg kg^−1^, subcutaneous injection) and post-operative meloxicam (1 mg/kg, subcutaneous injection). Mice were placed in a stereotaxic frame, their scalp hair removed, and the incision site sterilized using betadine and 70% ethanol. The skull was exposed and the conjunctive tissue removed using hydrogen peroxide. The skull was positioned such that the lambda and bregma marks were aligned on the anteroposterior and dorsoventral axes. For all surgeries, anti-inflammatory (Meloxicam) injections were pursued for 3 days following surgery.

For virus delivery, we first drilled a small craniotomy (0.5 mm) above the region of interest. For delivering virus in the mPFC or PPC, we injected a volume of 300–400 nl of virus (rate: 200 nl/min), using a glass pipette with a 50 μm diameter tip. Coordinates for targeting the mPFC/PPC virally were (in mm): anterior-posterior (AP) +1.5 to +2.0/-1.0, medial-lateral (ML): ±0.4/±1.5, dorsal-ventral (DV) − -1.75 and -2.25/ -0.5. We defined mPFC based on previous literature that included anterior cingulate cortex, prelimbic, and infralimbic cortex as part of PFC in rodents^51^. All injections were performed using an infuser system (QSI 53311, Stoelting) attached to the stereotaxic frame.

To deliver light into the mPFC or PPC, 200-μm two-ferrule cannulas (1.25mm ceramic ferrule 200 μm, Thorlabs, CFMLC12U-20) were implanted bilaterally above the mPFC or PPC using the following coordinates (in mm): mPFC: AP +1.5; ML: ±0.4; DV 1.85 at 10° in the ML axis; or PPC: AP: -2.0; ML: ±1.5; DV 0.4 at a 0° in the AP/ML axis. After implantation, dental cement (Teets Denture Material) and Metabond (C&B Metabond, Parkell) was applied to affix the implant to the skull. To avoid light reflection and absorption, the transparent Metabond was mixed with black ink pigment (Black Iron Oxide 18727, Schmincke). A custom designed head-plate was then positioned over the implant and affixed to the skull using Metabond. We used single ferrule cannulas with large (200 μm) diameter and high numerical aperture (0.39 NA) (Thorlabs, M89L01).

To perform mPFC and PPC single unit recording or optogenetic inhibition in awake head-fixed mice, we implanted a head plate parallel to the bregma–lambda axis of the skull. We used a custom design stereotactic arm to align the head plate parallel to the median and dorsal line of the skull during implantation. The head plate was attached to the skull using dental cement. The exposed skull was protected using rapid curing silicone elastomer (Kwik-Cast, WPI) topped with a fine layer of dental cement.

### Behavioral setup

Mice were head-fixed on a behavior rig and confined in a 3d-printed tube to limit body movements. A metallic lick spout position was controlled by mounting the spout apparatus on the 3d-printed stage controlled by miniature servo motor (Miuzei Mini Servo, MS18, China). The lick spout connected to a custom-made lick detector and was used to deliver water rewards (∼10 μl drop of water). A small tube, pointing toward the mouse facial area and at a distance of 3 cm, was used to deliver air-puff punishment (compressed air at 40 psi for 0.3 s). Voltage signals from the transducer and lick detector were recorded through a microcontroller board (Arduino UNO Rev3). A second microcontroller board was used to control a 5 mm white LED light placed 8 cm in front of the mouse right eye, and two solenoid valves (Parker 003-0141-900) for water and air-puff delivery. Three or fourteen kilohertz sound stimuli of 0.3 s duration were delivered using a single speaker located at a distance of 30 cm from the mouse. The speaker frequency range was calibrated using a USB calibrated measurement microphone (UMIK-1, Mini DSP) and the Room EQ Wizard software. We used five behavior rigs (four for general behavior, optogenetics, and training, one for electrophysiological recording). The behavioral setup was connected to a computer running a custom-written MATLAB (Mathworks) script that was able to detect licks while controlling the timing of the light cue, sound (using custom-written MATLAB code), lick port position, water, and reward. Behavior rigs were assembled primarily with optomechanical components (Thorlabs).

### Behavioral task and training

Upon recovery from the surgical procedure, mice were gradually put on a water restriction schedule, receiving eventually 1–1.6 ml of water in total per day. Body weight was maintained above 90% of the pre-restriction weight.

A white LED indicated the beginning of each trial. After a 1-1.5s delay, two auditory stimuli (0.3s duration) separated by a first-delay epoch (1.0 s) were presented and followed by another second-delay epoch (1.0 s). After waiting until the delay 2 epoch, the lick spout was rapidly moved within reach of the tongue, and remained within reach for 1.0 s if licks were not detected. Mice learned to lick the spout when they heard two identical tones (matched trial, 3-3/14-14 kHz frequencies in first- and second-stimulus) and to hold still when they heard two different tones (non-matched trial, 3-14/14-3 kHz). Correct licks during this period were rewarded with ∼10 μl drop of water and the spout remained within reach for an additional 2.0 s for licking water. Licks to the non-matched trial were punished with a siren-like auditory stimulus alone (early training) or siren plus air-puff (late training) and an extra 7.0 s inter-trial interval. At the end of the response epoch, the spout was then rapidly retracted and remained out of reach until the next trial (3-4s inter-trial interval).

Mice were taken through the following two stages of training until they became proficient at the task.

1. During the first phase of training (1st stage), mice learned to associate a lick with reward and to detect half category of matched/non-matched trials. In this phase, only 3-3 and 3-14 kHz pairs of tones were used. The delay 2 was set to 0.2 s and air-puff was not presented in this stage. Once mice performed successfully over 70% of trials in consecutive two days, they proceeded next stage.
2. During the second phase of training (2^nd^ stage), all pairs of tones were applied pseudo-randomly. The delay 2 was initially set to 0.2 s and air-puff was delivered only in false alarm trials. To prevent behavioral preference, an incorrect response resulted in the repetition of the same trial, thereby specifically increasing the trial length of the trial types with weak performance. Once they performed over about 100 trials, an incorrect trial was not repeated (test session). During the session, the performance correct rate (referred to as “correct rate” or “behavioral performance” in labels of figures) of each session was defined by: *Correct rate =* 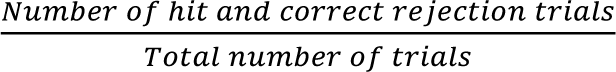. Once mice performed successfully over 70% (65% for optogenetic inactivation batches) of trials in two consecutive days, the second-delay was prolonged by 0.1 s additionally in the next session.

### Optogenetic inhibition of mPFC and PPC activity

Optical stimulation was applied through a ferrule-terminated 200 μm core and 0.39 NA optic fibre attached to the 200 μm core and 0.39NA patch cable (Thorlabs, M89L01) using a 1.25mm ceramic mating sleeve (Thorlabs, ADAL1-5). We used a blue-fibre-coupled light emission diode (λ = 470 nm, Thorlabs, M470F3). The light was delivered at 20Hz with a 0.4 duty cycle at an irradiance of 10mW mm^-2^ at the output tip of the fibre.

We performed optical stimulation at one of six timings (0-0.3s, 0.4-0.7s, 0.8-1.1s, 1.3-1.6s, 1.7-2.0s, or 2.1-2.4s relative to first-stimulus onset) pseudo-randomly. Optical stimulation trials were randomly interleaved two times out of every ten trials (20% of trials), and were not applied in two consecutive trials. The optical stimulation was delivered to both hemisphere/cannulas simultaneously.

### Electrophysiology

For in-vivo electrophysiology recordings, expert mice were anesthetized with isoflurane (3% induction, 1–1.5% for maintenance). They underwent a 1mm craniotomy (centered at 1.5/-2.0 mm anterior to bregma and ±0.4/1.5 mm lateral to the midline for mPFC/PPC). The dura was punctured and the craniotomy was protected with saline and a piece of gel form (Pfizer). The skull was covered again with silicone (Kwik-Cast, WPI) and the mouse was allowed to recover for at least 24h for the anesthesia effect to wash out completely.

The awake animal was then head-fixed and the silicone and gel form removed gently. 0.9% NaCl solution was used to keep the surface of the brain wet for the duration of the recordings. After placing the animal in the recording setup, we submerged a reference silver wire in the saline solution on the skull surface. The position of the NeuroPixels 1.0 probe^31^ (IMEC, Belgium) was referenced on the surface of the brain. The probe was then lowered slowly (about 1 min per mm), by manually insertion using a micromanipulator (MP-285, Sutter Instrument Company), until a depth of 3.0 mm was reached. Electrophysiological data and time stamps of each trial start were collected at a rate of 30kHz with a PXI based system (National Instruments), and saved using OpenEphys software (Neuropixels PXI plugin, Open ephys, https://open-ephys.github.io/gui-docs/User-Manual/Plugins/Neuropixels-PXI.html). We recorded up to 6 sessions per mouse. Probes were dipped in CM-DiI, DiD, or DiO (Invitrogen, ThermoFisher, #V22889) prior to insertion. In each session, we inserted up to 2 probes at a time. The probes were always inserted at the same angle within the coronal plane (-10° and 10° relative to the vertical axis for mPFC and PPC) to aid subsequent histological probe tract tracing.

Spike sorting was done using Kilosirt2.5^52^, and spikes were manually curated using Phy GUI (https://github.com/kwikteam/phy) to remove artifacts. Units with an inter-spike interval (ISI) violation ≤ 0.5, firing rate ≤ 0.1 Hz, or ill-shaped waveform were filtered out. Spike times were verified with cross correlograms to combine units or eliminate duplicates. For each unit, part of the recordings with obvious drift (unit spikes abruptly disappearing) were excluded.

### Histology

All mice were deeply anesthetized and transcardially perfused with 0.1 M phosphate-buffered saline (PBS) followed by 4% paraformaldehyde (PFA). Brains were dissected and post-fixed in 4% PFA overnight at 4°C. Brains were then sectioned into 100 μm coronal sections using a vibratome (Leica VT 1200S) and mounted on SuperFrost Plus slides (VWR) with Vectashield mounting media (Vector Laboratories). Brain sections were imaged with a confocal microscope (Leica SP8) using a 10x objective to confirm probe and implant positions in target regions.

### Analysis of behavior

To quantify behavior, we calculated hit and false alarm rates defined as follows:

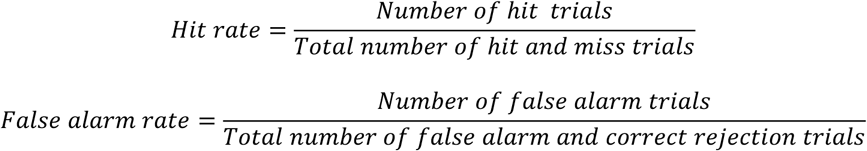

Discriminability (d’) was also used to quantify mice performance, as defined by:

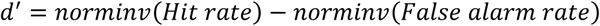

where, *norminv* was the inverse of the cumulative normal function.

For behavioral data obtained from optogenetic inactivation experiments (Extended Data Fig.1i), all behavioral metrics were calculated by pooling trials separately for mice injected with virus in the mPFC (n=3) and PPC (n=3), respectively. Statistical significance was assessed using a bootstrap test, performed by randomly sampling 50% of trials 1,000 times. Distributions were considered significantly dissociated if they showed no overlap within a 2SD range (containing 95.5% of data).

To quantify the impact of task and behavioral variables on behavior, we carried out regression analysis to weigh the contributions of stimulus and history of correctness, stimulus, false alarm, hit, and choice on the animal’s choice on the current trial^53^ (Extended Data Fig.1e-f). To do so, we concatenated data from multiple sessions for each mouse and fit the animal’s choice with a logistic regression model as follows:

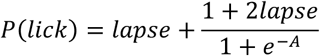

where

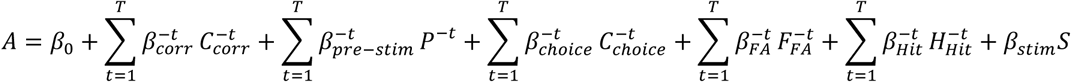

where 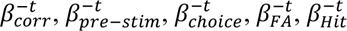 and *β_stim_* are coefficients of the 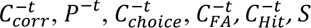, regressors, respectively. *T* is the number of trials in the past. For this model, we took into account for up to 3-trial history. In this equation (5), 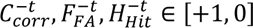 if a trial is correct or incorrect, false alarm or not, and hit or not at *t*-back trial, *P*^−*t*^ ∈ [+1, −1] if a trial is a matched or non-matched at *t*-back trial, 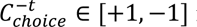 if the mice licked or non-licked a port at *t*-back trial, *S* ∈ [+1, −1] if the current trial is a matched or non-matched, respectively. *β*_0_ is the intercept of the model that captures the overall bias of mice. We used the negative log-likelihood as the cost function *J*:

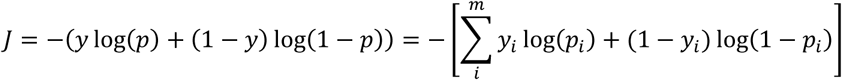

where *m* is the total number of trials. The model was fit using a gradient-descent algorithm to minimize the negative log-likelihood cost function with elastic net regularization with parameters *α* = 0.2 (L1) and *λ* = 0.8 (L2). We used the sqp algorithm in the *fmincon* function from MATLAB. To compare the impact of parameters of current stimulus and previous trial information, we also fit the model without current stimulus (*S*) or previous trial parameters (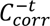, 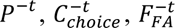, and 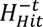). All models were fit using 200 runs of fivefold cross-validation. For each run, we computed the log-likelihood for the test dataset given the best-fit parameters on the training set (log*l*). We also calculated the log-likelihood of the test dataset given the only best-fit parameters of *β*_0_. This gives us a null log-likelihood reference value (log*l*_0_). In order to quantify the efficiency of each model we defined the cross-validated bit/trial (CV-bit/trial) as the trial-averaged excess likelihood of the model compared to the null model^53^:

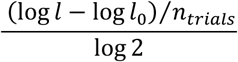

To compare different models, we calculated the mean value of CV-bit/trial across 200 runs for each mouse.

### Analysis of neural activity

For all mPFC and PPC neurons, changes in firing rate associated with behavior were assessed using peri-stimulus time histogram (PSTHs). Unless otherwise stated, PSTHs were computed using 10 ms bins for individual neurons in each recording session and smoothed with a Gaussian filter with a standard deviation of 20 ms to obtain the temporal profile. For visualization and analysis, firing rates were z-scored across sessions.

#### Selectivity

All comparative indices (stimulus 1, 2, and choice selectivity index) were computed using a receiver operating characteristic (ROC) analysis (Fig.1h and Extended Data Fig.2), which calculates the ability of an ideal observer to classify whether a given spike density was recorded under one of two trial types. We indexed the difference between two firing rate distributions by scaling the ROC area between -1 and 1, where 0 reflects no difference between the distributions and the sign denotes whether a neuron fires more under one of the trial types than the other. Statistical significance (p<0.05) was determined with a permutation test of 500 repetitions. In this analysis, we used firing rates in only correct trials during the task.

We assessed stimulus 1 and 2 selectivity by comparing high and low tone trials at the stimulus (0-0.3s and 1.3-1.6s relative to stimulus 1 onset) or delay period (0.8-1.3s and 2.1-2.6s relative to stimulus 1 onset). Choice selectivity was computed by comparing lick and non-lick trials at the delay period.

#### Bootstrap test

We evaluated the statistical significance in the analysis using data resampling with a bootstrapping procedure. We estimated the *p*-value for the bootstrapping procedure by computing the ratio (1 + *X*) / (*N* + 1), where *X* indicates overlapping data points between the two distributions, and *N* indicates iterations.

#### State-space analysis

For state-space analysis (Extended Data Fig.3), we used neurons that were recorded in the sessions with >65% correct rate. To characterize the population structure and temporal pattern among all neurons during the analysis window (stimulus subspace: 0-0.5s and 1.3-1.8s relative to stimulus 1 onset, memory maintenance subspace: 0.8-1.3s and 2.1-2.6s), z-scored firing rates were formatted as *X* ∈ R*^N^*^×*CT*^, where *N* is the number of neurons, *C* is the total number of conditions (e.g., high/low tones in stimulus 1 and 2 period), and *T* is the number of analyzed time points. Principal component analysis (PCA) was used to reduce the dimensionality of the population from the number of neurons to ten principal components (PCs). Each PC represents a weighted combination of individual neuronal activity, which summarizes population activity.

To statistically estimate the difference between each neuronal trajectory at each time point across trial types in the PC space, we computed each PC using a subset of trials (50%) and projected the data from the remaining subset of trials (50%) onto each axis. The projection of each axis was computed by the dot product as *v_pc_x*, where *v_pc_* is a weight vector for each PC, and *x* is an *N* × (4 × *time*) matrix of smoothed, trial-averaged firing rates across trial types (low-low, low-high, high-low, and high-high). The procedure was repeated 100 times with shuffling trials within each trial type. We then computed the Euclidean distance between low and high tone trials (stimulus) or lick and non-lick trials (choice) using the first 3 or 10 dimensions. Statistical significance was tested by comparing the resampling distributions of distance between empirical and shuffled-label data. If the distributions were not overlapped in a 2SD range (95.5% data in this range), they were defined as significantly dissociated.

#### Linear regression model

We used a linear regression model to determine how various task variables affect the responses of each neuron (Supplementary Fig.3). We first z-scored the responses of a given neuron by subtracting the mean response from the firing rate at each time and in each trial and by dividing the result by the standard deviation of the responses. Both the mean and the standard deviation were computed by combining the neurons responses across all trials and times. We then described the z-scored responses of neuron *i* at time *t* as a linear combination of several task variables (Supplementary Fig.3a):

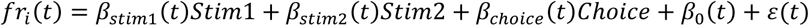

where *fr_i_*(*t*) is the z-scored response of neuron *i* at time *t*, *Stim*1 indicates the category of stimulus 1 (+1: low tone; -1: high tone), *Stim*2 indicates the category of stimulus 2 (+1: low tone; -1: high tone), and *C*ℎ*oice* represents the animal’s choice (+1: licked; 0: non-licked). *β*_0_(*t*) and *εε*(*t*) denote the intercept and residual of the z-scored response, respectively.

To estimate the optimal weights for each variable (*β_stim_*_1_(*t*), *β_stim_*_2_(*t*), *β_c_*_ℎ*oice*_ (*t*), and *β*_0_(*t*)) while preventing overfitting, the models were fitted to each neuron’s response using the *lassoglm* function in MATLAB with five-fold cross validation of the training data. We specified a normal distribution with a linear (identity) link function and set the hyperparameter *λ* to 0.01. Only neurons recorded in the sessions with >65% correct rate were included in the linear regression model analysis.

For assessing the significance of pairwise correlations, we computed Pearson’s correlation coefficients (*r*) and their associated *p*-values using the *corr* function in MATLAB. The function calculates two-tailed p-values under the null hypothesis of no correlation (*r* = 0) by transforming the correlation coefficient into a t-statistic.

#### Classification analysis

For classification analysis (Fig.2a-b, 3c, 3f, Extended Data Fig.4a-b, 4e, 5a-b, 5d-f, 5i, 8i, Supplementary Fig.1b-c, Supplementary Fig.2a, 2d, and Supplementary Fig.4a, 4c), we used neurons that were recorded in the sessions with >65% correct rate. For classifiers, we used a logistic regression classifier as implemented by the MATLAB *fitclinear* function to quantify the amount of information about the stimulus (low or high tone) and choice (lick or non-lick trial) in the population of neurons recorded from each brain region. This analysis included all neurons that were recorded during at least 40 trials for each stimulus and choice trial types, and we constructed “pseudo-trials” by randomly extracting trials from desired conditions for each neuron^54^. We used only correct trials for the classification analysis. For the training and testing dataset, the number of trials in each condition was matched to prevent bias in training classifiers. We also randomly chose the trials of lick trials from low-low and high-high trials and non-lick trials from low-high and high-low trials with equal probability (20 trials for each combination). Classifiers with L2 regularization (λ = 1/20) was trained to classify the stimulus or choice using pseudo-population data and tested on held-out data. The choice of the λ value did not affect our conclusion. We used tenfold cross-validation by leaving a 10% subset of trials for classification to avoid overfitting. This procedure was repeated 100 times. Classification results are reported either as a function of time or in a fixed time window. Time-resolved classification was done on z-scored firing rates measured in a 50 ms moving window. For fixed-window classification, we used z-scored firing rates in a 500ms for stimulus (0-0.5s or 1.3-1.8s from stimulus onset) and delay (0.8-1.3s or 2.1-2.6s from stimulus onset) periods. For cross-conditional classification approach, we trained a classifier to classify one condition (e.g., high/low tone trials) using a pseudo-population data from the epoch/time corresponding to the condition (e.g., delay 1 epoch), and tested on held-out data from another epoch/time corresponding to another condition (lick/non-lick in delay 2 epoch). Statistical significance was tested by comparing the resampling distribution of classification accuracy averaged within tenfold cross-validation against chance level (chance level = 0.5 for stimulus and choice classification). If the distribution and the chance level are not overlapped in a 2SD range (95.5% data in this range), they are defined as significantly dissociated. Statistical tests were performed independently for mPFC and PPC because they represent separate datasets and distinct priori hypotheses; therefore, no correction for multiple comparisons between mPFC and PPC was applied.

#### Angle between subspaces

For calculation of angle between axes (Fig.2c, Fig.3d, 3e, 3g, 3h, Extended Data Fig.4c, 4f, 5c, 5g, and Supplementary Fig.2b, 2e), the decision boundary of a logistic regression classifier can be computed as

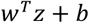

where *w* is the weights, *z* is z-scored firing rates, and *b* is an intercept. We obtained *w^cond^*^1^ and *w^cond^*^2^ for each condition and computed angle *θ* between these as:

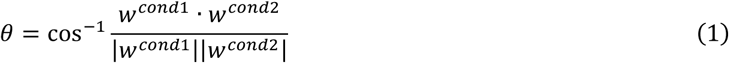

The angle *θ* was converted from radian to degree ranged 0 to 90 degrees. Statistical significance was tested by comparing the resampling distribution of empirical angles and angles computed based on the weights obtained from classifiers trained with shuffled-label data.

For calculation of angle between axes for each animal (Fig.3e, 3h), we first calculated the condition-averaged z-scored firing rate (high/low, or lick/non-lick) for each neuron for the relevant epoch. To obtain the “Coding Direction^55,56^”, we computed the difference of firing rates between conditions. Specifically, we defined the following two axes for this analysis. The “stimulus axis” was computed from the dissociation of neuronal activity between the low and high tone trials in the stimulus period (0-0.5s, or 1.3-1.8s from first-stimulus onset). The “memory maintenance axis” was computed from dissociation of neuronal activity between low and high tone trials or lick and non-lick trials in the latter-half of delay period (0.8-1.3s, or 2.1-2.6s from first-stimulus onset). We prepared the four vectors, which are average population activity *v* of length *N_unit_* × 1. Each vector was normalized by their unit length as *v_norm_* = *v*⁄‖*v*‖. The angle between stimulus axes was calculated as equation (1) using *v_norm_* calculated using stimulus periods. The angle between memory maintenance axes was calculated using *v_norm_* from memory periods.

To establish the statistical significance of the observed Spearman rank correlation coefficient (*C_obs_*) between behavioral performance and the angle of subspaces, a two-tailed permutation test was performed. A null distribution was generated by randomly shuffling the angle relative to the behavioral performance and calculating the correlation coefficient for each iteration (*C_perm_*). This procedure was performed 1000 times. The final empirical P-value was calculated as the proportion of absolute correlation coefficients from the shuffled distribution that were greater than or equal to the absolute value of the observed correlation as follows:

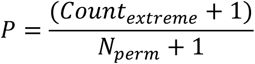

where *Count_extreme_* is the number of permuted correlations satisfying |*C_perm_*| ≥ |*C_obs_*|and *N_perm_* is the number of permutations. (Fig.3e, h, and Extended Data Fig.5f, P<0.05, Spearman rank-correlation test).

#### Geometric analysis

For geometric analysis (Fig.3i-j), we used neurons that were recorded in the sessions with >65% correct rate. To visualize the geometry of trials in the population activity, z-scored firing rates were formatted as *X* ∈ ℝ*^N^*^×8^, where *N* is the number of neurons and four combinations of tones (low-low, low-high, high-low, and high-high tones) at two stimulus periods (0-0.5, and 1.3-1.8s from stimulus 1 onset) or the latter-half of delay periods (0.8-1.3, and 1.8-2.3s from stimulus 1 onset). PCA was used to reduce the dimensionality of the population from the number of neurons to a lower number of principal components (PCs).

For the geometric analyses within each session (Extended Data Fig.7c-e, 7f-g), we used sessions with behavioral performance above 65% and at least five simultaneously recorded neurons. Following the same procedure described above, the population activity was reduced from the number of recorded neurons to a lower-dimensional space defined by PCs. The number of PCs was chosen such that they collectively explained more than 70% of the total variance. Within this reduced space, we defined coding directions as vectors connecting the mean population activity between specific trial conditions. Specifically, the coding direction during the late delay 1 epoch was defined from low-tone trials (L–L and L–H) to high tone trials (H-L and H-H), and that during the late delay 2 epoch was defined from Go trials (L–L and H–H) to No-go trials (L–H and H–L). Each coding direction was represented as a vector in Euclidean space connecting the mean positions of the corresponding trial conditions. To quantify whether the two coding directions align with similar or opposite directions, we calculated the cosine similarity between them. Positive values indicate acute angles (i.e., similar directions), while negative values indicate obtuse angles (i.e., opposite directions). For each session, we randomly resampled each unique trial 10 times and computed the cosine similarity. This procedure was repeated 100 times to avoid sampling bias.

To study how the geometry of the first-two PC space evolved throughout the trial, the projections of the first-two axes were computed as the same procedure described in state-space analysis at each time throughout the trial (Extended Data Fig.6a-d).

#### Clustering analysis

To examine the dominance of specific clusters of neurons contributing to the stimulus and memory maintenance subspace, we conducted clustering analysis based on neurons’ response profile to the task variables (Fig.4c-g). First, we assume that we have recorded a response profile *R* ∈ ℝ*^N^*^×*q*^, where *N* is the number of neurons and *q* is the number of features. The features were defined as the average firing rate during four different epochs (0-0.5s, 0.8-1.3s, 1.3-1.8s, and 2.1-2.6s from stimulus 1 onset) for each unique combination of tones, resulting in 16 features in total (Fig.4a). The response profile matrix *R* was then normalized (z-scored) for each neuron. Second, we applied the GMM to the normalized response profile matrix *R*. To do that, we used the *fitgmdist* function in MATLAB with a 0.35 regularization value, 100 replicates, and the covariance matrix constrained to diagonal^57^.

To determine the number of clusters, we computed the Bayesian information criteria (BIC) score (Fig.4e). It is a penalized likelihood term defined as 2(*N* log *L*) + *M* log *n*, where *N* log *L* is the negative log-likelihood of the data, *M* is the number of parameters of the GMM and *n* is the number of observations. The BIC score was computed by the *fitgmdist* function.

#### ePAIRS

To statistically test whether mPFC and PPC neurons exhibited clusters of prototypical response profiles or a uniform continuum of response profiles, we carried out the ePAIRS statistical test^24,42^ (Fig.4b, 4d, Extended Data Fig.8a), which is itself derived from the PAIRS test developed in ^22^. We used a response profile *R* described above as a representational space and it was z-scored for each neuron. For each neuron, we computed the median vector angle *α_i_* with its *k*-nearest neighbors (*k* being a hyperparameter set to 3 in this study), defining an empirical distribution 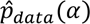. For comparison, we generated null distributions that exhibited no clustering. A multivariate Gaussian distribution N(0, ∑) is fit to the response profile *R* using *mvnrnd* function in MATLAB, with ∑ being the empirical covariance of *R*, computed as 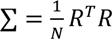. We then computed the median vector angle of nearest neighbors for this simulation dataset, defining a null distribution *p^_null_*(*α*). The difference between the *p^_data_* and *p^_null_* is assessed using a two-sided Wilcoxon rank-sum test^42^.

#### Cell-type classification

To classify the putative fast-spiking (FS) interneuron and regular-spiking (RS) excitatory neurons, we calculated trough-to-late peak and peak-to-valley ratio for each recorded neuron (Extended Data Fig.8f). Neurons were clustered into two groups using spectral clustering with a correlation-based distance metric and a random-walk normalized Laplacian. Following clustering, we computed the mean firing rate of neurons within each cluster and assigned cluster identities as putative FS or RS neurons based on the higher and lower firing rates, respectively.

### RNNs

We built a rate-based RNN where each unit’s activity matched the PSTH of an experimentally recorded neuron from the mPFC, pooled across sessions^58,59^ (Fig.6). All networks are a time-discretized RNN with positive activity. Before time discretization, the network state ℎ evolves according to a dynamical equation:

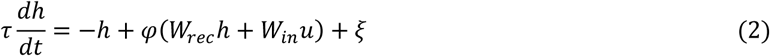

Here, *τ* is the units’ time constant. *φ*(·) is the non-linear activation function, the sigmoid function was used in this study, W*_rec_* and W*_in_* are the recurrent and input weight matrices, *u* is the inputs to the network, and *ξ* represents Gaussian noise.

The network’s output units *z* was read out from the network according to:

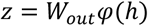

where W*_out_* is the output connection matrix of size *N_out_* × *N_rec_*. All weight matrices (W*_in_*, W*_rec_*, W*_out_*) were learned over the course of training.

After using the first-order Euler approximation with a time-discretization step Δ*t*, we have

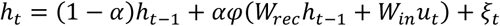

Here, *α* ≡ Δ*t*/*τ*, and we use a discretization step Δ*t* = 20ms. We imposed no constraint on the sign or the structure of the weight matrices W*_in_*, W*_rec_*, W*_out_*. The network and the training are implemented in PyTorch.

The network received three types of noisy input:

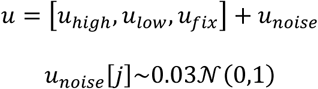

The fixation input *u_fix_* was 1 when the network was required to fixate and 0 when the network was required to respond. The stimulus inputs (*u*_ℎ*iii*ℎ_ and *u_loll_*) represented high and low tones in the mouse’s experiment, respectively, and were set to 1 during the stimulus epoch and 0 otherwise.

#### Task and performance

We implemented an abstraction of the DMS-dr task that mice performed in this study (Fig.6a). The task consisted of six periods; initial delay (100 ms), stimulus 1 (300 ms), delay 1 (1000 ms), stimulus 2 (300 ms), delay 2 (1000 ms), and response period (200 ms). Inputs and outputs for an example trial are shown in Fig.5a. Fixation input remained 1 throughout the trial until the response period, where it changed to 0. If the target fixation output prematurely fell below 0.5, the network was deemed to have erroneously broken fixation, resulting in an incorrect trial. Fixation input and output reflect the suppression of licking behavior in the mouse’s experiment. Stimulus inputs represented high and low tones in the experiment. The average of response output during the response period determined whether the trial was reported as a match (*output* > 0.5) or non-match (*output* ≤ 0.5), reflecting licking or non-licking behavior in the mouse’s experiment. The task performance was computed by comparing the response output with successfully reported with match and non-match trials.

#### Network training procedure

We used back-propagation through time^60^ to train networks to minimize loss functions ℒ. Trials were generated stochastically, following a predefined temporal structure and mapping stimulus inputs to target outputs *z^*. The total loss function was:

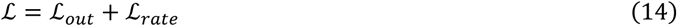

where

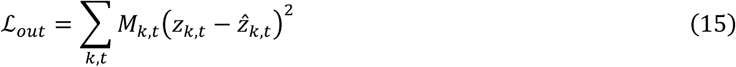

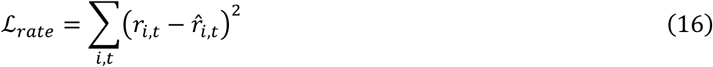

Here, *z_k_*_,*t*_ and *z^_k_*_,*t*_ are, respectively, the actual and the target readout values and the indices *k* and *t*, respectively, are the index of the output and timesteps. *r_i_*_,*t*_ and *r^_i_*_,*t*_ are, respectively, the actual and the trial-averaged mPFC neuronal activity of the *i*-th neuron. We implemented a mask, *M_k_*_,*t*_, for modulating the loss with respect to certain time intervals.

- For the response output unit (*k* = 1), *M*_1,*t*_ = 0 before stimulus 1 onset, *M*_1,*t*_ = 5 during the response period, and *M*_1,*t*_ = 1 for the rest of the trial.
- For the fixation output unit (*k* = 2), *M*_2,*t*_ = 0 before stimulus 1 onset and *M*_2,*t*_ = 2 for all other timepoints.

We trained networks with *N* = 312 and *N* = 77 units to reproduce neural activity recorded from mPFC clusters 2 and 3 (Fig.4a), respectively. Additionally, we included other 61 neurons which were not trained to reproduce neuronal activity recorded from the mPFC. The total number of units in the network was *N* = 450. The networks were trained using Adam optimizer^61^ in PyTorch with the decay rate for the first and second moment estimates of 0.9 and 0.999 and learning rates of 10^−2^, respectively. During training, we used mini-batches of 32 trials. In a given minibatch, all trial types were randomly generated. Training was terminated when ℒ*_out_* fell below 0.02, and the task performance exceeded over 0.99.

#### Unit inactivations

We silenced a network unit by setting its activity to zero for all specific cluster of neurons at given timepoints. Units were silenced at five epochs: initial delay (0-100ms from trial start), stimulus 1 (100-500ms), late-delay1 (900-1400ms), stimulus 2 (1400-1800ms), and late-delay 2 (2200-2700ms) (Fig.6c).

#### Network simulation of training history

To investigate how training history influences representational geometry in the network, we simulated learning of the DMS-dr task RNNs (Extended Data Fig. 7a-e). For each condition, we trained 10 independent RNNs consisting of 128 recurrent units with a sigmoid activation function.

- Pre-training: Networks were first pre-trained on simplified versions of the DMS-dr task containing only two trial types. Specifically, “Low-associated” networks were trained on L–L and L–H trials, whereas “High-associated” networks were trained on H–H and H–L trials. Training was terminated once the output loss (ℒ*_out_*) fell below 0.005.
- DMS-dr task training: Following pre-training, networks were trained on the full DMS-dr task containing all four trial types (L–L, L–H, H–L, and H–H), using the same convergence criterion (ℒ*_out_* < 0.005). In addition, a separate cohort of networks was trained on the DMS-dr task without any pre-training.

To quantify the degree to which memory-related subspaces were shared, we analyzed population activity from trained RNNs during task performance. For each network, recurrent unit activity was averaged across trials for each condition and segmented into delay 1 (0.8–1.3 s from stimulus 1 onset) and delay 2 (2.1–2.6 s) epochs. Delay-period activity across all conditions was z-scored for each unit. PCA was applied to this activity matrix. We retained the minimum number of principal components required to explain at least 90% of the variance. Population activity was projected onto this low-dimensional subspace, and coding directions were defined as vectors connecting condition centroids. A stimulus coding direction was defined by the difference between Low and High tone conditions, and a memory-related coding direction was defined by the difference between matched and non-matched trials. The cosine similarity between these two vectors was computed to quantify the alignment between two memory representations, following the same geometric analysis described above. Cosine similarity values were computed for each network and compared across three cohorts: Low-associated networks, High-associated networks, and networks trained without pre-training. Statistical significance was assessed using one-sample t-tests against zero.

## Statistics and reproducibility

Statistical tests and sample sizes are reported in the figure captions. No statistical methods were used to predetermine sample sizes. Animals were not randomly assigned to experimental groups, and investigators were not blinded to group allocation during experiments or outcome assessment. The locations of Neuropixels probes and optical fiber implants were confirmed histologically in all animals. Sessions in which animals did not perform the behavioral task, yielded too few recorded neurons, or contained too few trials were excluded from analysis. Detailed exclusion criteria are described in the Methods section. In optogenetic experiments, mice lacking detectable viral expression were excluded.

Statistical analyses were performed using MATLAB. Unless otherwise stated, all statistical tests were two-sided, and P < 0.05 was considered statistically significant. Data were compared using t-tests or analysis of variance (ANOVA) followed by pairwise comparisons with multiple-comparison correction using the Tukey–Kramer method. For normally distributed data, pairwise comparisons were performed using t-tests with Holm–Bonferroni correction. For classification analyses, significance was assessed using one-sided permutation tests. Classification performance is presented as the mean ± SD, corresponding to approximately 95.5% of observations under a normality assumption.

## Data availability

Data to generate the figures are available via Mendeley Data at https://doi.org/10.17632/h92tsjrwg5.1.

## Code availability

All code supporting the main findings from this study are available via Mendeley Data at https://doi.org/10.17632/h92tsjrwg5.1.

## References

1. Koechlin, E., Ody, C. & Kouneiher, F. The architecture of cognitive control in the human prefrontal cortex. Science 302, 1181–1185 (2003).

2. Badre, D. & D’Esposito, M. Functional magnetic resonance imaging evidence for a hierarchical organization of the prefrontal cortex. J. Cogn. Neurosci. 19, 2082–2099 (2007).

3. Badre, D. & Nee, D. E. Frontal cortex and the hierarchical control of behavior. Trends Cogn. Sci. 22, 170– 188 (2018).

4. Driscoll, L. N., Shenoy, K. & Sussillo, D. Flexible multitask computation in recurrent networks utilizes shared dynamical motifs. Nat. Neurosci. 27, 1349–1363 (2024).

5. Yang, G. R., Joglekar, M. R., Song, H. F., Newsome, W. T. & Wang, X.-J. Task representations in neural networks trained to perform many cognitive tasks. Nat. Neurosci. 22, 297–306 (2019).

6. Cole, M. W. Cognitive flexibility as the shifting of brain network flows by flexible neural representations. Current Opinion in Behavioral Sciences 57, 101384 (2024).

7. Stanley, M. L., Gessell, B. & De Brigard, F. Network Modularity as a Foundation for Neural Reuse. Philos. Sci. 86, 23–46 (2019).

8. Fu, Z. et al. The geometry of domain-general performance monitoring in the human medial frontal cortex. Science 376, eabm9922 (2022).

9. Minxha, J., Adolphs, R., Fusi, S., Mamelak, A. N. & Rutishauser, U. Flexible recruitment of memory-based choice representations by the human medial frontal cortex. Science 368, eaba3313 (2020).

10. Willett, F. R. et al. Hand knob area of premotor cortex represents the whole body in a compositional way. Cell 181, 396–409.e26 (2020).

11. Courellis, H. S. et al. Abstract representations emerge in human hippocampal neurons during inference. Nature 632, 841–849 (2024).

12. Panichello, M. F. & Buschman, T. J. Shared mechanisms underlie the control of working memory and attention. Nature 592, 601–605 (2021).

13. Bernardi, S. et al. The Geometry of Abstraction in the Hippocampus and Prefrontal Cortex. Cell 183, 954–967.e21 (2020).

14. Tafazoli, S., et al. Building compositional tasks with shared neural subspaces. bioRxiv (2024) doi:10.1101/2024.01.31.578263.

15. Jahn, C. I. et al. Learning attentional templates for value-based decision-making. Cell 187, 1476–1489.e21 (2024).

16. Boyle, L. M., Posani, L., Irfan, S., Siegelbaum, S. A. & Fusi, S. Tuned geometries of hippocampal representations meet the computational demands of social memory. Neuron 112, 1358–1371.e9 (2024).

17. Nogueira, R., Rodgers, C. C., Bruno, R. M. & Fusi, S. The geometry of cortical representations of touch in rodents. Nat. Neurosci. 1–12 (2023).

18. Romo, R., Brody, C. D., Hernández, A. & Lemus, L. Neuronal correlates of parametric working memory in the prefrontal cortex. Nature 399, 470–473 (1999).

19. Machens, C. K., Romo, R. & Brody, C. D. Flexible control of mutual inhibition: a neural model of two-interval discrimination. Science 307, 1121–1124 (2005).

20. Liu, D. et al. Medial prefrontal activity during delay period contributes to learning of a working memory task. Science 346, 458–463 (2014).

21. Stroud, J. P., Watanabe, K., Suzuki, T., Stokes, M. G. & Lengyel, M. Optimal information loading into working memory explains dynamic coding in the prefrontal cortex. Proc. Natl. Acad. Sci. U. S. A. 120, e2307991120 (2023).

22. Raposo, D., Kaufman, M. T. & Churchland, A. K. A category-free neural population supports evolving demands during decision-making. Nat. Neurosci. 17, 1784–1792 (2014).

23. Churchland, M. M. et al. Neural population dynamics during reaching. Nature 487, 51–56 (2012).

24. Hirokawa, J., Vaughan, A., Masset, P., Ott, T. & Kepecs, A. Frontal cortex neuron types categorically encode single decision variables. Nature 576, 446–451 (2019).

25. Yang, W., Tipparaju, S. L., Chen, G. & Li, N. Thalamus-driven functional populations in frontal cortex support decision-making. Nat. Neurosci. 1–14 (2022).

26. Wu, Z. et al. Context-Dependent Decision Making in a Premotor Circuit. Neuron 106, 316–328.e6 (2020).

27. Yu, L. et al. The causal role of auditory cortex in auditory working memory. Elife 10, e64457 (2021).

28. Marton, T. F., Seifikar, H., Luongo, F. J., Lee, A. T. & Sohal, V. S. Roles of Prefrontal Cortex and Mediodorsal Thalamus in Task Engagement and Behavioral Flexibility. J. Neurosci. 38, 2569–2578 (2018).

29. Wilhelm, M. et al. Striatum-projecting prefrontal cortex neurons support working memory maintenance. Nat. Commun. 14, 7016 (2023).

30. Baeg, E. H. et al. Dynamics of population code for working memory in the prefrontal cortex. Neuron 40, 177–188 (2003).

31. Jun, J. J. et al. Fully integrated silicon probes for high-density recording of neural activity. Nature 551, 232–236 (2017).

32. Meyers, E. M., Freedman, D. J., Kreiman, G., Miller, E. K. & Poggio, T. Dynamic population coding of category information in inferior temporal and prefrontal cortex. J. Neurophysiol. 100, 1407–1419 (2008).

33. Stokes, M. G. et al. Dynamic coding for cognitive control in prefrontal cortex. Neuron 78, 364–375 (2013).

34. Libby, A. & Buschman, T. J. Rotational dynamics reduce interference between sensory and memory representations. Nat. Neurosci. 24, 715–726 (2021).

35. Mante, V., Sussillo, D., Shenoy, K. V. & Newsome, W. T. Context-dependent computation by recurrent dynamics in prefrontal cortex. Nature 503, 78–84 (2013).

36. Maaten, L. & Hinton, G. E. Visualizing Data using t-SNE. Journal of Machine Learning Research 9, 2579– 2605 (2008).

37. Osako, Y. et al. Contribution of non-sensory neurons in visual cortical areas to visually guided decisions in the rat. Curr. Biol. 31, 2757–2769.e6 (2021).

38. Fascianelli, V. et al. Neural representational geometries reflect behavioral differences in monkeys and recurrent neural networks. Nat. Commun. 15, 1–19 (2024).

39. Bellafard, A., Namvar, G., Kao, J. C., Vaziri, A. & Golshani, P. Volatile working memory representations crystallize with practice. Nature 629, 1109–1117 (2024).

40. Komiyama, T. et al. Learning-related fine-scale specificity imaged in motor cortex circuits of behaving mice. Nature 464, 1182–1186 (2010).

41. Posani, L., Wang, S., Muscinelli, S. P., Paninski, L. & Fusi, S. Rarely categorical, always high-dimensional: how the neural code changes along the cortical hierarchy. bioRxiv 2024.11.15.623878 (2025) doi:10.1101/2024.11.15.623878.

42. Dubreuil, A., Valente, A., Beiran, M., Mastrogiuseppe, F. & Ostojic, S. The role of population structure in computations through neural dynamics. Nat. Neurosci. 1–12 (2022).

43. Zatka-Haas, P., Steinmetz, N. A., Carandini, M. & Harris, K. D. Sensory coding and causal impact of mouse cortex in a visual decision. bioRxiv 501627 (2021) doi:10.1101/501627.

44. Guo, Z. V. et al. Flow of cortical activity underlying a tactile decision in mice. Neuron 81, 179–194 (2014).

45. Pho, G. N., Goard, M. J., Woodson, J., Crawford, B. & Sur, M. Task-dependent representations of stimulus and choice in mouse parietal cortex. Nat. Commun. 9, 2596 (2018).

46. Vinograd, A., Nair, A., Kim, J. H., Linderman, S. W. & Anderson, D. J. Causal evidence of a line attractor encoding an affective state. Nature 634, 910–918 (2024).

47. Forli, A. et al. Two-photon bidirectional control and imaging of neuronal excitability with high spatial resolution in vivo. Cell Rep. 22, 3087–3098 (2018).

48. Carrillo-Reid, L., Han, S., Yang, W., Akrouh, A. & Yuste, R. Controlling Visually Guided Behavior by Holographic Recalling of Cortical Ensembles. Cell 178, 447–457.e5 (2019).

49. Daie, K., Svoboda, K. & Druckmann, S. Targeted photostimulation uncovers circuit motifs supporting short-term memory. Nat. Neurosci. 24, 259–265 (2021).

50. Marshel, J. H. et al. Cortical layer-specific critical dynamics triggering perception. Science 365, eaaw5202 (2019).

51. Anastasiades, P. G. & Carter, A. G. Circuit organization of the rodent medial prefrontal cortex. Trends Neurosci. 44, 550–563 (2021).

## Methods only references

52. Pachitariu, M., Sridhar, S., Pennington, J. & Stringer, C. Spike sorting with Kilosort4. Nat. Methods 21, 914– 921 (2024).

53. Akrami, A., Kopec, C. D., Diamond, M. E. & Brody, C. D. Posterior parietal cortex represents sensory history and mediates its effects on behaviour. Nature 554, 368–372 (2018).

54. Rigotti, M. et al. The importance of mixed selectivity in complex cognitive tasks. Nature 497, 585–590 (2013).

55. Allen, W. E. et al. Thirst regulates motivated behavior through modulation of brainwide neural population dynamics. Science 364, 253 (2019).

56. Li, N., Daie, K., Svoboda, K. & Druckmann, S. Robust neuronal dynamics in premotor cortex during motor planning. Nature 532, 459–464 (2016).

57. Engelhard, B. et al. Specialized coding of sensory, motor and cognitive variables in VTA dopamine neurons. Nature 570, 509–513 (2019).

58. Rajan, K., Harvey, C. D. & Tank, D. W. Recurrent Network Models of Sequence Generation and Memory. Neuron 90, 128–142 (2016).

59. Finkelstein, A. et al. Attractor dynamics gate cortical information flow during decision-making. Nat. Neurosci. 24, 843–850 (2021).

60. Werbos, P. J. Backpropagation through time: what it does and how to do it. Proc. IEEE Inst. Electr. Electron. Eng. 78, 1550–1560 (1990).

61. Kingma, D. P. & Ba, J. Adam: A method for stochastic optimization. arXiv [cs.LG] (2014).

