## Supplementary figures and images for "Reusable modular architecture enables flexible cognitive operations in the mouse brain and artificial recurrent networks"

### Extended Data Fig.1

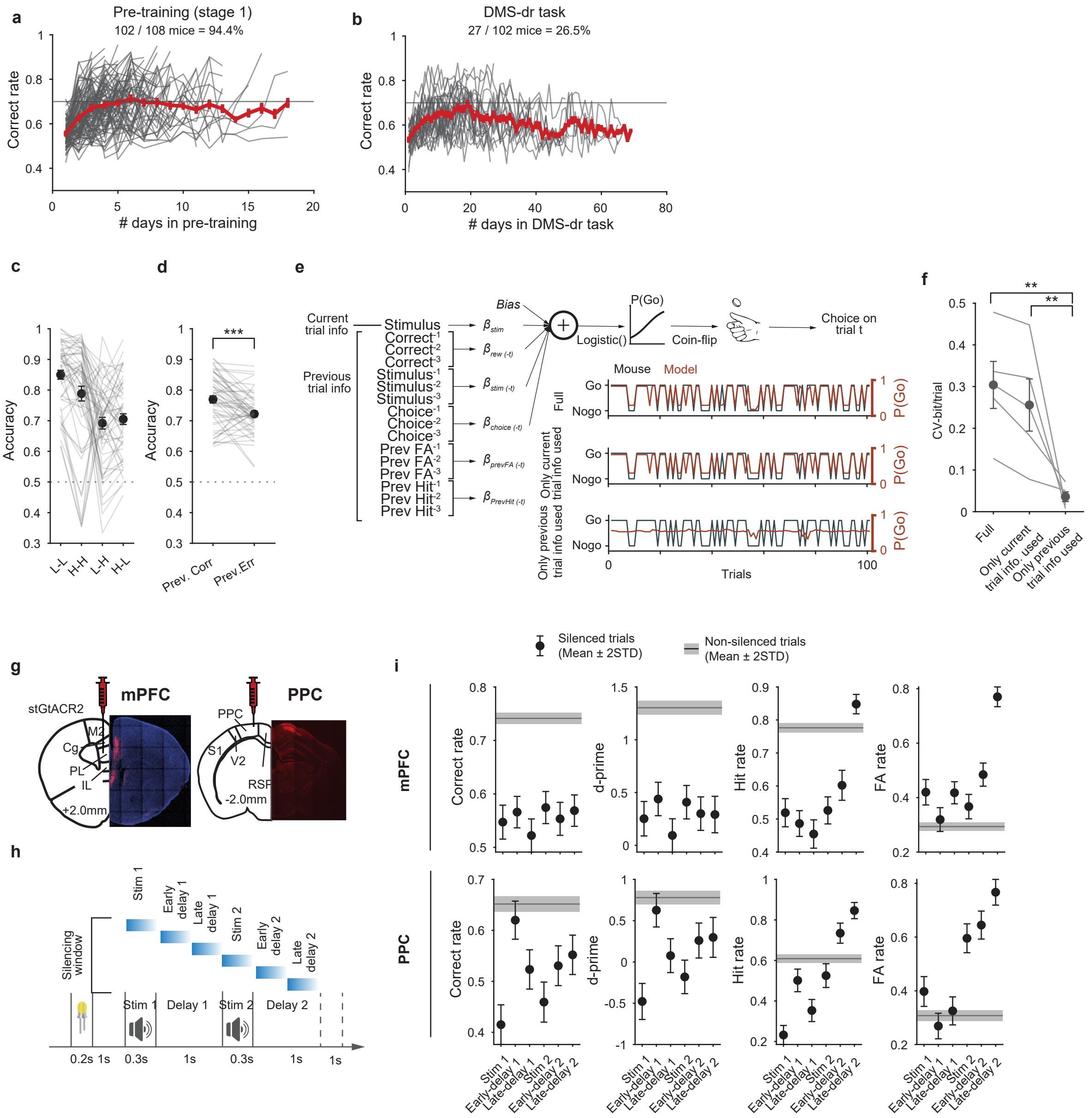

### Extended Data Fig.2

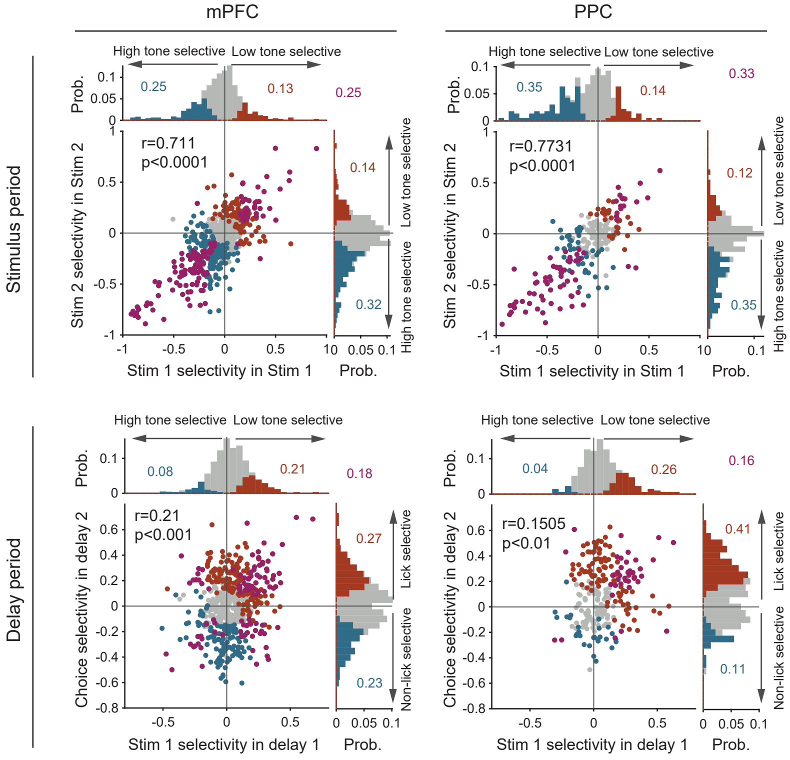

### Extended Data Fig.3

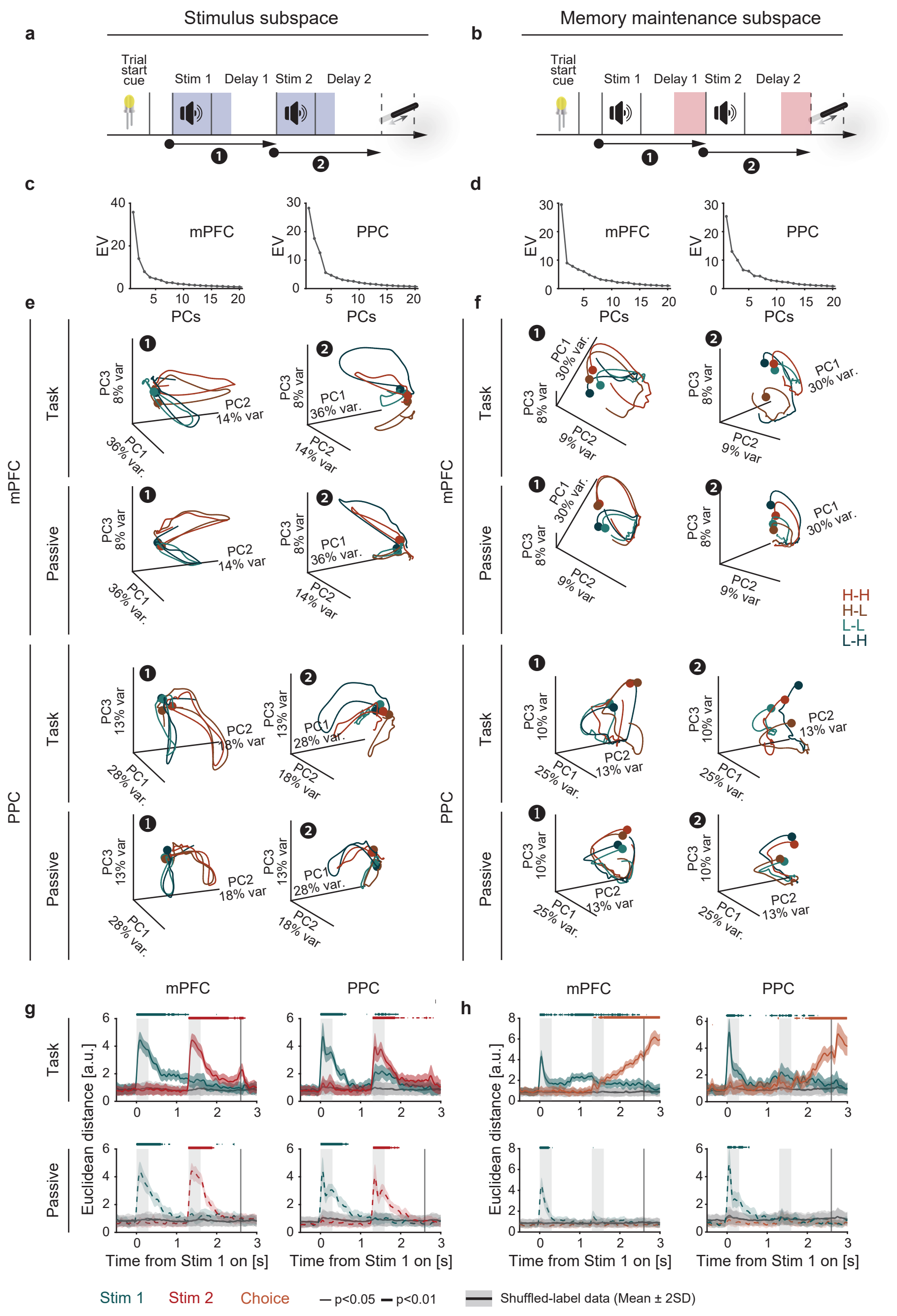

### Extended Data Fig.4

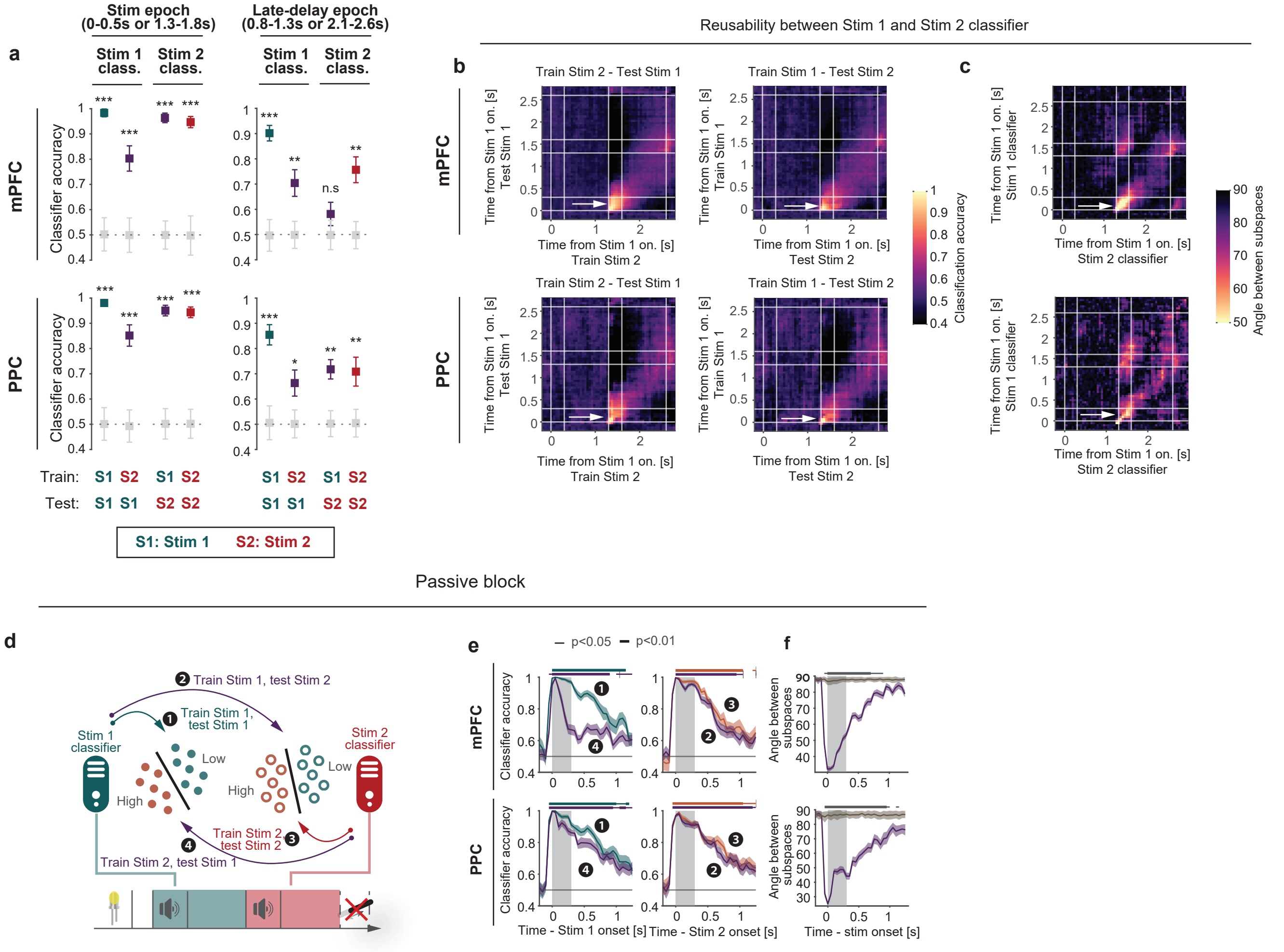

### Extended Data Fig.5

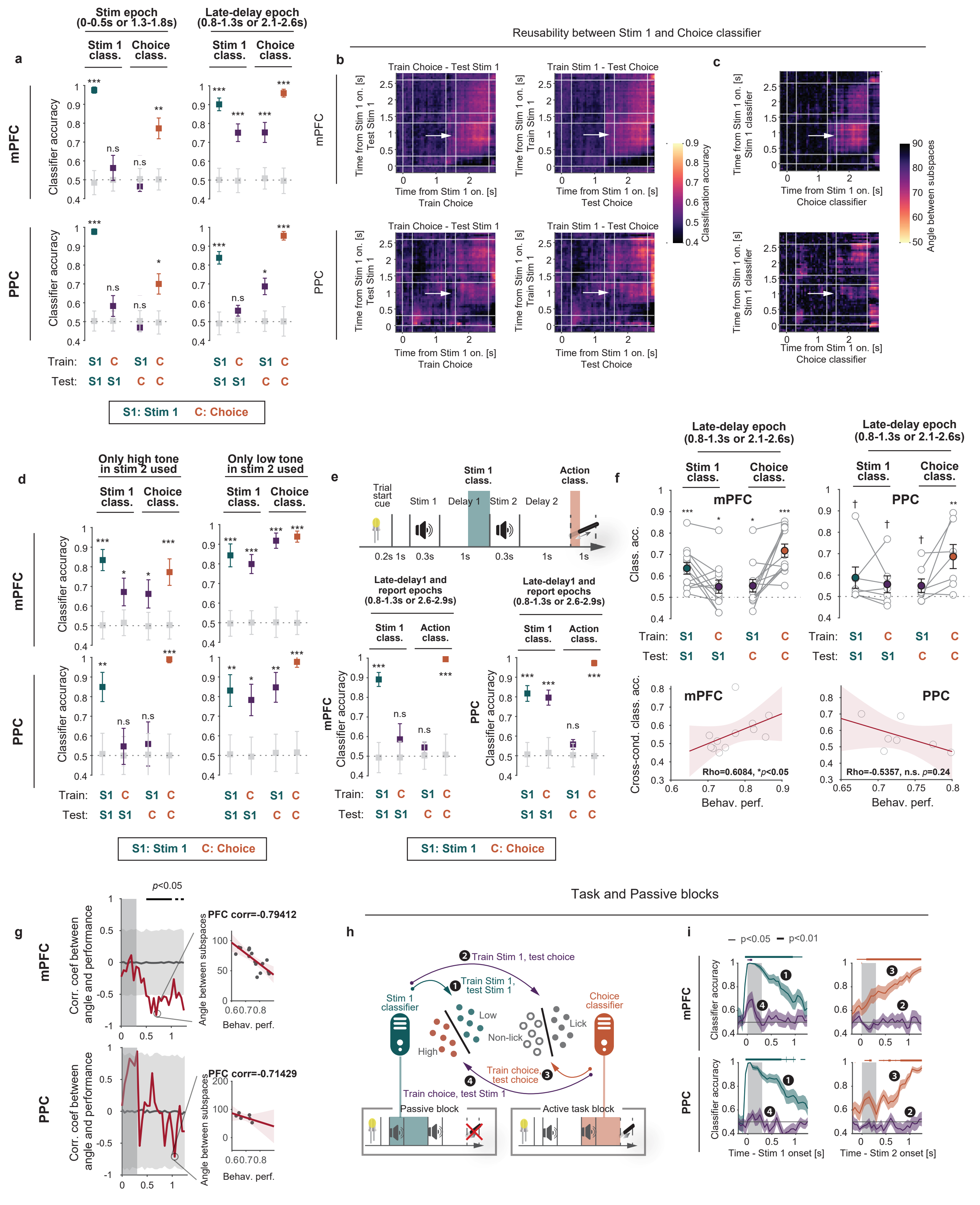

### Extended Data Fig.6

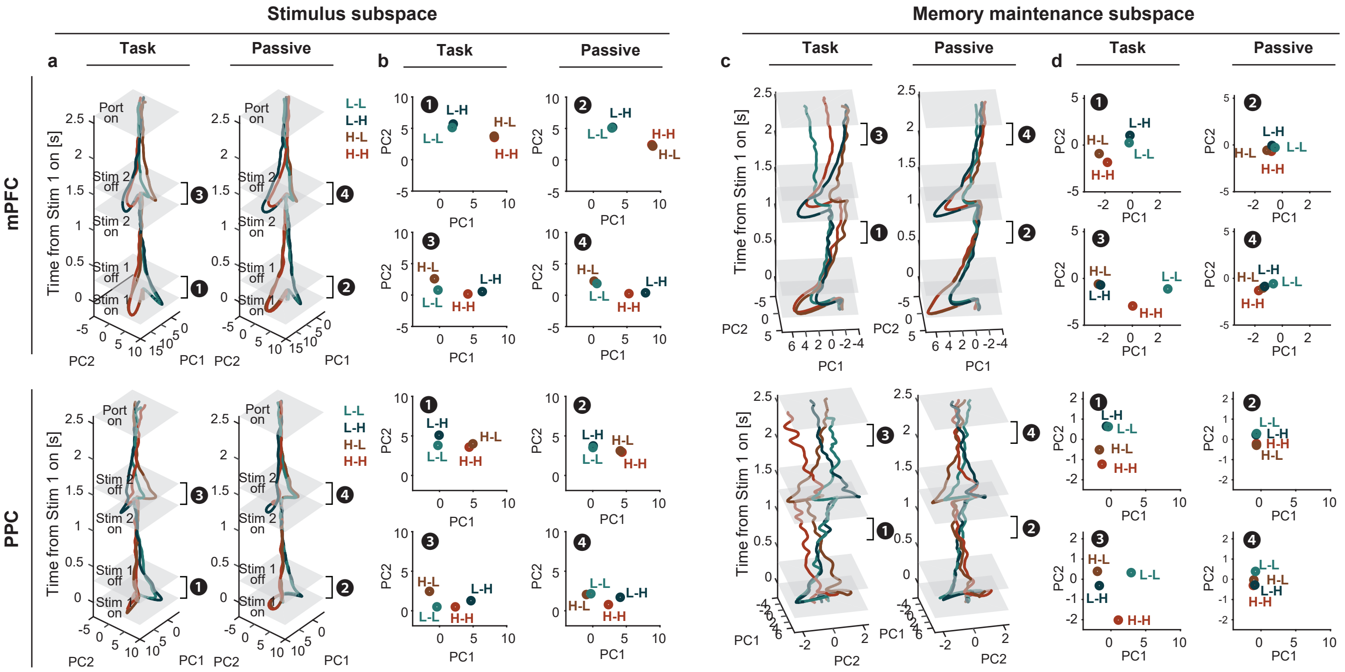

### Extended Data Fig.7

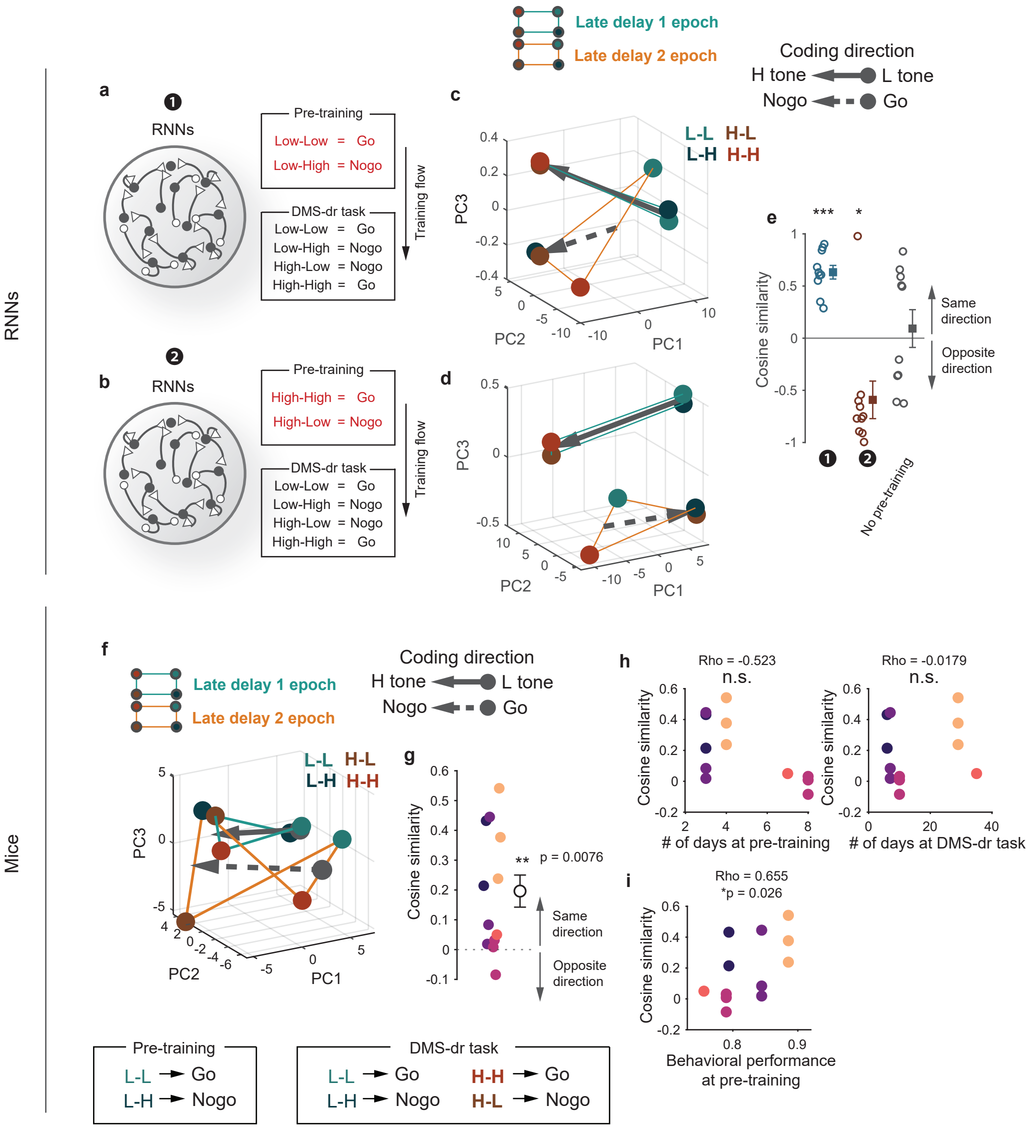

### Extended Data Fig.8

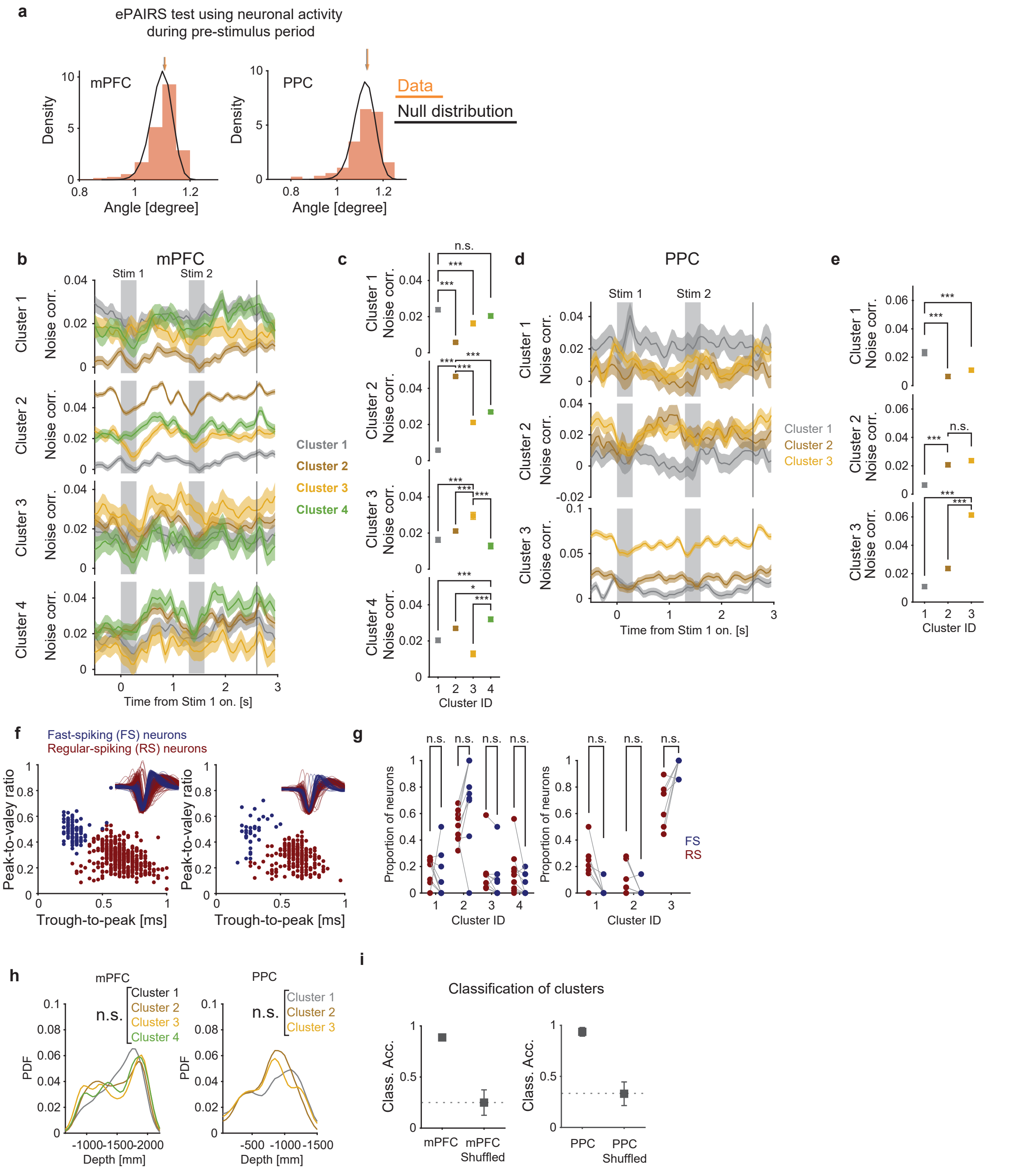

### Extended Data Fig.9

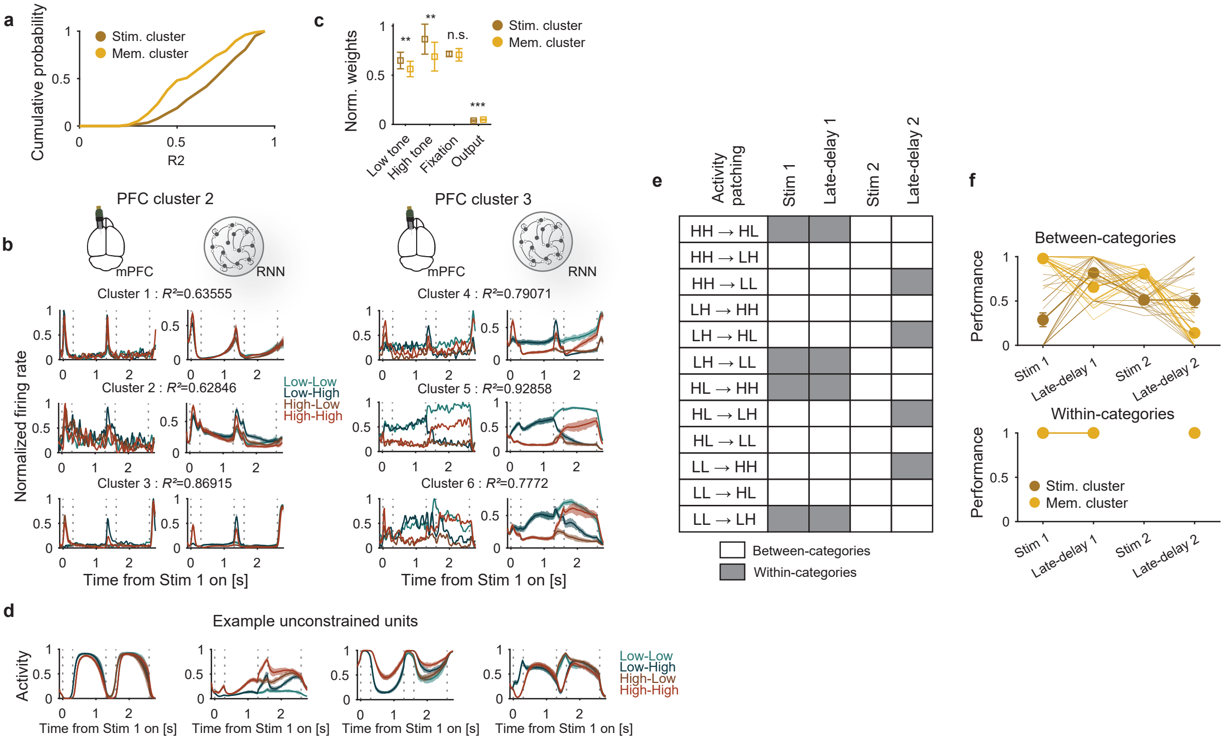
